# TLR5 drives metabolic dysfunction-associated steatohepatitis through lipid- and flagellin-induced hepatocyte injury signalling

**DOI:** 10.64898/2026.02.05.703969

**Authors:** Wenhao Li, Niannian Wang, Raju Kumar, Iris Gines Mir, Upkar Gill, Gillian Hood, James Brindley, Charles Mein, James Boot, Rose Wilcox, Neil Dufton, Robert Goldin, John Loy, Kalpana Devalia, Humza Malik, Adam Goralcyzk, Maria Jimenez Ramos, Timothy J. Kendall, Jonathan Fallowfield, Maria Ines Castanho Martins, Krista Rombouts, Michele Vacca, Dania El Abyad, Seray Anak, Olivier Govaere, William Alazawi

**Affiliations:** Barts Liver Centre, Blizard Institute, Queen Mary University of London, London, UK; Genome Centre, Blizard Institute, Queen Mary University of London, London, UK; William Harvey Research Institute, Queen Mary University London, London, UK; Department of Metabolism, Digestion and Reproduction, Imperial College London, London, UK; Homerton University Hospital NHS Foundation Trust, London, UK; Institute for Regeneration and Repair, University of Edinburgh, Edinburgh, UK; Regenerative Medicine and Fibrosis Group, Institute for Liver and Digestive Health, Royal Free Hospital, University College London, London, UK; Clinica Medica “Frugoni”, Interdisciplinary Department of Medicine, University of Bari “Aldo Moro”, Bari, Italy; Roger Williams Institute of Liver Studies, School of Immunology & Microbial Sciences, Faculty of Life Sciences and Medicine, King’s College London, Foundation for Liver Research and King’s College Hospital, London, UK; Department of Imaging and Pathology, Translational Cell and Tissue Research, University Hospitals Leuven, KU Leuven, Leuven, Belgium

**Author notes:** Corresponding author: Professor William Alazawi Barts Liver Centre, Blizard Institute, Queen Mary University of London, London, UK.

## Abstract

Liver fibrosis is a strong predictor of clinical outcomes in metabolic dysfunction-associated steatohepatitis (MASH). Fibrosis is a consequence of persistent liver cell injury and inflammation in which Toll-like receptors (TLRs) play a key initiating role. Here we test the hypothesis that TLR5 is involved in the development of MASH and fibrosis using a combination of clinical data from multiple independent patient cohorts, single cell liver transcriptomics and human in vitro and ex vivo models. Hepatic TLR5 expression, but not TLR2 or TLR4, is associated with liver fibrosis and mortality. Plasma levels of TLR5’s cognate ligand flagellin are increased in MASH with advanced fibrosis and fall with liver disease improvement. Mechanistically, we identify two parallel TLR5-mediated routes to hepatocyte injury: one elicited by flagellin and the other indirectly by lipid injury. Furthermore, hepatocyte TLR5 inhibition abrogates paracrine activation of hepatic stellate cells to suppress collagen production. This is also seen ex vivo in patient-derived precision-cut liver slices where TLR5 inhibition significantly reduces lipid-induced collagen deposition. These findings reveal a new role for TLR5 signalling, specifically in the development of advanced MASH fibrosis and may offer a novel disease-specific therapeutic approach.

---

The degree of liver fibrosis predicts liver-related outcomes in people living with metabolic dysfunction-associated steatotic liver disease (MASLD) and its progressive form, metabolic dysfunction-associated steatohepatitis (MASH). MASH fibrosis can lead to cirrhosis, liver failure, and cancer ^1^ ^2^ ^3^. A complex interplay of metabolic factors results in excess lipid accumulation in the liver. In some patients, this leads to cellular stress responses, inflammatory signalling and production of cytokines which activate fibrogenic pathways in a range of cells, including hepatic stellate cells (HSC).

The source of the drivers of damage pathways is incompletely understood, but a prevailing hypothesis is that the gut–liver axis plays a critical role. Obesogenic diets contribute to intestinal dysbiosis ^4^, bacterial overgrowth ^5^, increased intestinal permeability and bacterial translocation ^6^, facilitating delivery of microbial products (pathogen-associated molecular patterns - PAMPs) and excess lipids to the liver ^7^ ^8^. In MASH, free fatty acids (FFAs) such as palmitic acid and oleic acid (most abundant saturated and unsaturated fats in the typical Western diet ^9^), accumulate in hepatocytes resulting in lipotoxicity. This leads to release of pro-inflammatory cytokines and damage-associated molecular patterns (DAMPs). PAMPs and DAMPs bind hepatic Toll-like receptors (TLRs), triggering pro-inflammatory cascades that promote liver injury and fibrogenesis.

Alongside lipid-mediated injury, there is an increased abundance of flagellated bacteria in MASH, mainly from the Proteobacteria and Firmicutes phyla which account for up to 50% of the human intestinal microbiome ^10^ ^11^ ^12^ ^13^. Flagellated bacteria express flagellin, the structural protein of bacterial appendages on the outer bacterial wall used for locomotion. Flagellin is the cognate ligand for TLR5 ^14^ which, when bound, activates MyD88-dependent NF-κB signalling, leading to production of inflammatory cytokines including interleukin(IL-)8 ^15^ ^16^. In vitro, IL-8 is released by hepatocytes following lipotoxic injury ^17^. Human plasma IL-8 concentrations are higher in individuals with MASH compared to those without and is independently associated with histological features of hepatocyte injury (ballooning) and fibrosis ^18^. We hypothesise that hepatic TLR5 signalling is activated in MASH and that modulating this pathway attenuates inflammatory signalling (IL-8 production) and subsequent fibrogenesis.

Previous murine TLR5 knockout/inhibition studies have demonstrated conflicting, model-dependent results ^19^ ^20^ ^21^. Global TLR5 knockout mice on high fat diet for 8 weeks ^19^ and hepatocyte-specific TLR5 knock out mice on methionine- and choline-deficient diet for 4 weeks^20^ demonstrated worsening liver steatosis and inflammation. However, TLR5 antagonism via inhibitor TH1020 improved liver steatosis, inflammation and fibrosis in C57BL/6N mice fed high-fat, high-fructose diet for 16 weeks ^21^. Furthermore, a meta-analysis by Vacca et al.^22^ demonstrated significant model-dependent variability in phenotype, liver transcriptome and histology across 39 commonly used murine models of MASLD. To mitigate against these inconsistencies, clinical and patient-relevant human cell and tissue studies are needed.

Here we show that hepatic TLR5 expression and circulating levels of flagellin are associated with clinically-relevant endpoints such as the stage of fibrosis, major liver events and mortality. Both human in vitro and ex vivo systems show that lipid uptake into hepatocytes (in the absence of flagellin) leads to a TLR5-dependent injury response that can activate HSC to drive fibrosis.

## Results

### Increased hepatic TLR5 expression in advanced MASH fibrosis is associated with liver-related events and mortality

TLR5 gene expression in whole-tissue liver biopsy mRNA was significantly greater in advanced MASH fibrosis group (fibrosis stages F3–F4) compared to control (n = 23, **Figure 1a, Supplementary Table 1**). TLR5 protein co-localised with albumin (**Figure 1b**), indicating hepatocyte expression. This is consistent with an independent published single nuclear RNAseq dataset ^23^ which showed 92.9% of TLR5 expression is from hepatocytes in human MASLD (**Supplementary Figure 1a**). TLR5 expression was predominantly observed outside fibrotic regions when analysed with spatial RNA sequencing (CosMx, **Figure 1c**) and did not co-localise with α-SMA **(Figure 1d)**, a marker of activated HSC.

**Figure 1.**
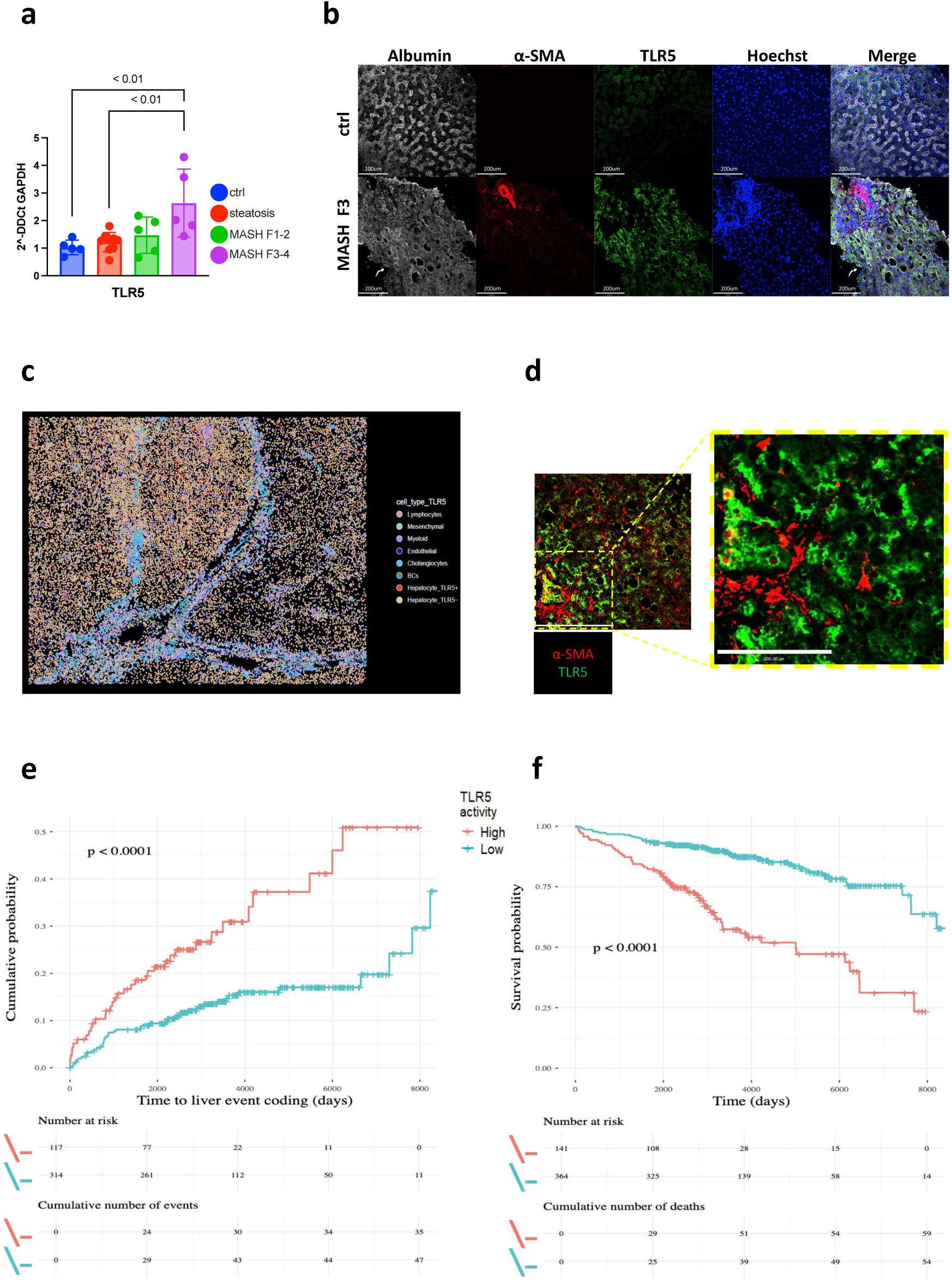
Hepatic TLR5 expression is increased in advanced MASH fibrosis and associated with adverse patient outcomes. **a**, Quantitative PCR of TLR5 from whole liver biopsy tissue, grouped by histological stage (ctrl n=5, steatosis n=9, MASH F1-2 n=5, MASH F3-4 n=5). **b**, Representative images of immunofluorescence of human liver biopsy tissue (n=12) showing TLR5 (green), albumin (grey), α-SMA (red), Hoescht (blue) in control (top row) and MASH with F3 fibrosis (bottom row) (Scale bar 200µm). **c**, NanoString CosMX analysis of explant liver tissue from MASH advanced fibrosis. Red dots represent TLR5 expressing cells. **d**, Zoomed representative images of immunofluorescence staining of human liver biopsy tissue showing TLR5 (green) and α-SMA (red) in MASH F3 (n=12,scale bar 100µm). **e**, **f**, Kaplan–Meier curves for liver-related events and all-cause mortality stratified by hepatic TLR5 expression in the SteatoSITE cohort. Error bars represent mean±standard deviation; P values determined by t test or ANOVA with post-test multiple comparisons. ANOVA, analysis of variance. Only statistically significant comparisons (p<0.05) are highlighted.

TLR5, but not TLR2 or TLR4, was upregulated in advanced MASH fibrosis in three other independent biopsy-proven MASLD cohorts when compared to early-stage disease or healthy controls (cohort 1 – SteatoSITE, n = 408; cohort 2 – combined University of Cambridge (UCAM), n = 58 and Virginia Commonwealth University (VCU), n = 78; cohort 3: Newcastle University (EPoS), n = 168; **Supplementary Figure 1b–d; Supplementary Table 2**). In the SteatoSITE cohort, high TLR5 expression was significantly associated with increased rates of liver-related events/hepatic decompensation (n = 431 with no liver event coding prior to biopsy, and death as a competing risk) and all-cause mortality (n = 505; Kaplan–Meier analysis, log-rank test, P<0.01 for both; **Figure 1e,f**). The association of high hepatic TLR5 expression with increased all-cause mortality remained significant in both early (F0-2) and advanced (F3-4) fibrosis stages (**Supplementary Figure 1e,f**). Taken together, these results show that hepatic (principally hepatocyte) TLR5 expression is increased in advanced MASH fibrosis and significantly associated with patient outcomes.

### Plasma flagellin increases in advanced MASH fibrosis and decreases with disease regression

The known ligands for TLR5 are the bacterial protein flagellin and the mammalian damage-associated protein High Mobility Group Box 1 (HMGB1) ^24^. Plasma flagellin, but not HMGB1 or gram-negative bacterial lipopolysaccharide (LPS), was increased in advanced MASH fibrosis compared to earlier disease stages (n = 61, **Figure 2a, b, Supplementary Figure 2a)**. Consistent with this, stool *FliC* gene abundance was higher in advanced MASH fibrosis group (n = 52, mean increased 28-fold, p=0.03; **Figure 2c**). Plasma flagellin did not change with worsening liver fibrosis in a non-obese cohort with treatment-naive chronic hepatitis B virus infection (n = 18, **Supplementary Figure 2b, Supplementary Table 3**), suggesting an increase in flagellin may be specific to MASLD. Furthermore, levels of plasma flagellin, but not LPS, decreased in MASLD participants following metabolic surgery compared to baseline (n = 36, mean reduction 1.4-fold, p<0.01; **Figure 2d, 2e**). Samples were taken pre-operatively on the day of surgery and a median of 89 days post-operatively when body weight and non-invasive markers of MASLD stage (controlled attenuation parameter and transient elastography liver stiffness) had also decreased (**Supplementary Figure 2c, d, e**). Collectively, findings from patient cohorts provide a rationale for deeper mechanistic investigation of TLR5 signalling in hepatocyte injury induced by flagellin, lipids (FFAs) or their combination.

**Figure 2.**
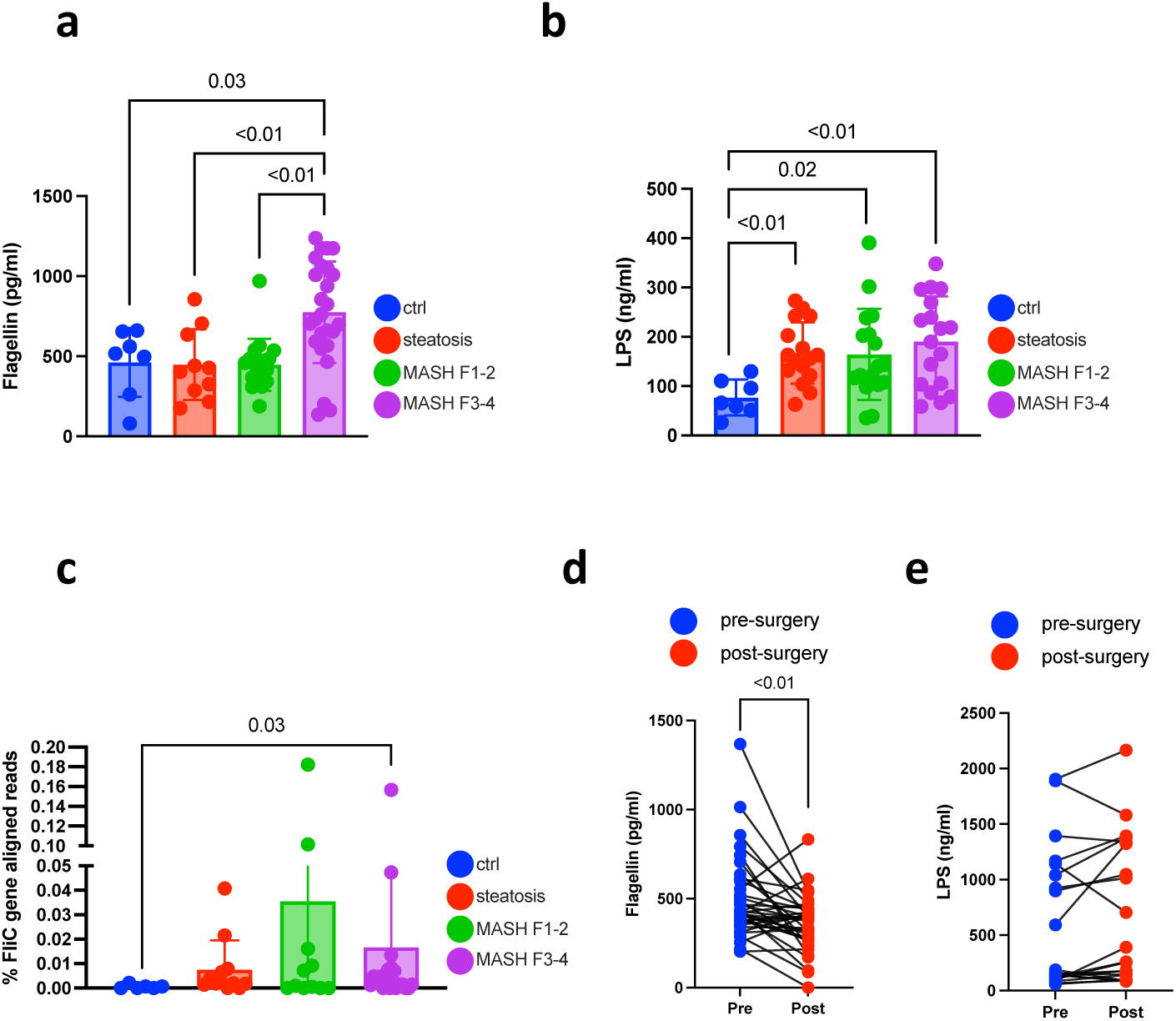
Circulating and microbial markers of TLR5–flagellin signalling in MASLD across fibrosis stages and following metabolic surgery. **a**, Plasma flagellin concentration in 61 participants by histological stage (control n=7, steatosis n=10, MASH F1-2 n=18, MASH F3-4 n=26). **b**, Plasma LPS concentration in 58 participants by histological stage (control n=7, steatosis n=16, MASH F1-2 n=17, MASH F3-4 n=18). **c**, Stool flagellin gene load expressed as *FliC* gene reads as percentage of total gene reads per sample extracted from stool DNA isolated from 52 participants by histological stage (control n=6, steatosis n=12, MASH F1-2 n=13, MASH F3-4 n=21). **d**, Paired changes in plasma flagellin concentration, and **e**, Paired changes in plasma LPS concentration following metabolic surgery in 36 biopsy-proven MASLD participants. Error bars represent mean ± standard deviation; P values determined by t test or ANOVA with post-test multiple comparisons. ANOVA, analysis of variance. Only statistically significant comparisons (p<0.05) are highlighted.

### TLR5 inhibition reduces flagellin-induced and lipid-induced IL-8 production in hepatocytes

In vitro, purified *Proteobacteria*-derived flagellin induced the production of pro-inflammatory IL-8 protein in primary hepatocytes (**Figure 3a**) and cell lines (HepG2, Huh7, **Supplementary Figure 3a,b**). IL-8 production was attenuated by inhibition with the selective TLR5 receptor antagonist, TH1020 ^25^ (**Figure 3a**). In line with in vitro findings, plasma IL-8 was elevated in participants with advanced MASH fibrosis compared to healthy controls (**Supplementary Figure 3c**) and was positively associated with histological markers of hepatocyte injury (NAFLD activity score) and fibrosis stage (**Supplementary Figure 3d,e**).

**Figure 3.**
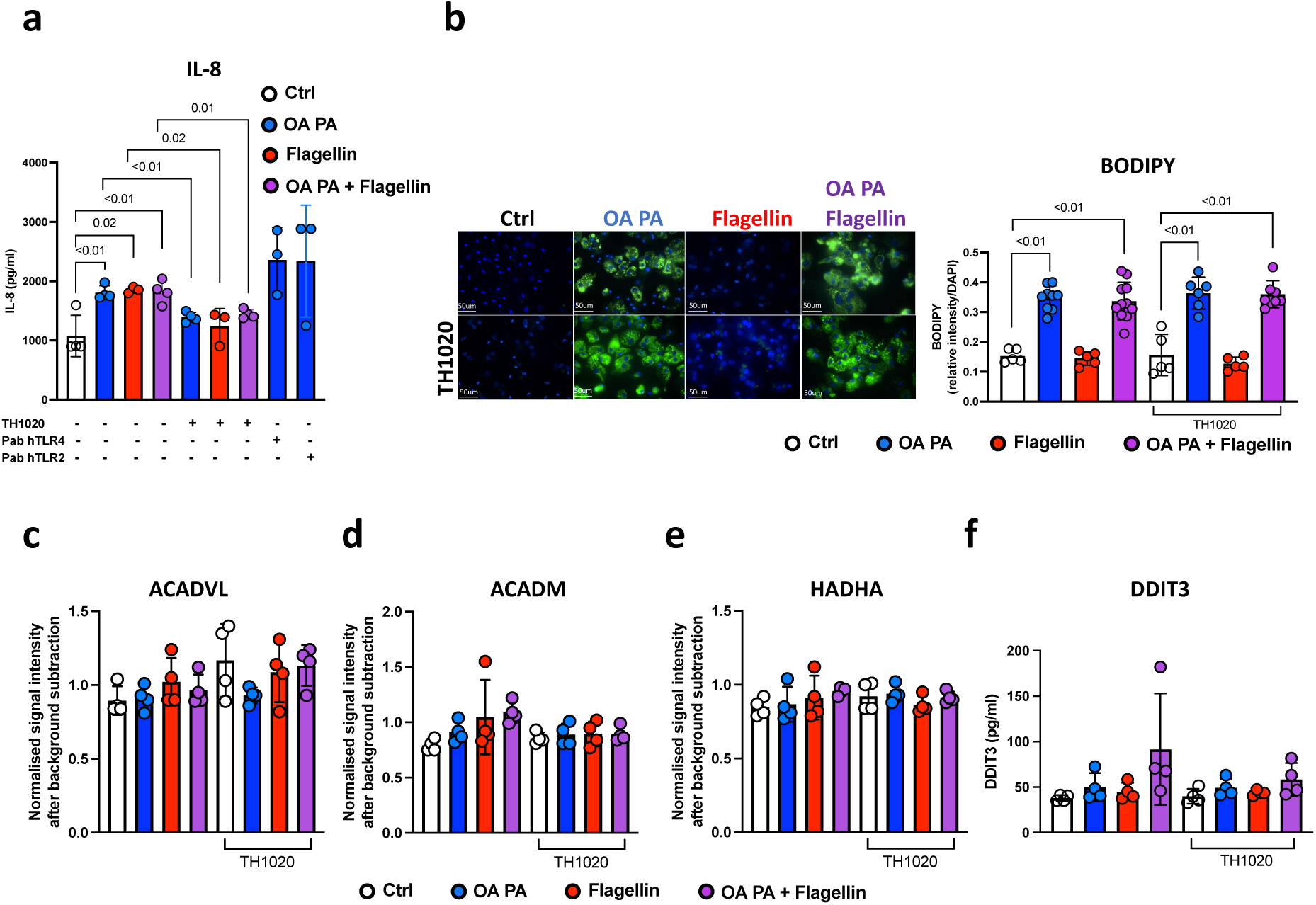
TLR5 inhibition reduces flagellin- and lipid-mediated IL-8 production in primary hepatocytes but does not affect lipid accumulation, expression of proteins associated with fatty acid oxidation or endoplasmic reticulum stress. Primary hepatocytes treated with oleic acid and palmitic acid (OA PA), flagellin or combination (OA PA + flagellin) for 24 hours **a,** in the presence or absence of 1µM TH1020 (TLR5 inhibitor) , 5µg/mL Pab hTLR4 (TLR4 inhibitor), 5µg/mL Pab hTLR2 (TLR2 inhibitor), **b** - **f** in the presence or absence of 1µM TH1020. **a**, IL-8 concentration (n=3–4 replicates per condition). **b**, Representative images of lipid deposition quantified using BODIPY 493/503 staining (green), nuclei stained with DAPI (blue) via confocal microscopy (Scale bars 50μm) and BODIPY quantified according to relative intensity/DAPI. Experiments performed in triplicates for each condition. **c - e** Expression of fatty acid oxidation enzymes: **c,** very long chain specific acyl-CoA dehydrogenase – ACADVL, **d,** medium-chain specific acyl-CoA dehydrogenase - ACADM and **e,** long-chain 3-hydroxyl-CoA dehydrogenase – HADHA. **f**, DNA damage-inducible transcript 3 (DDIT3) concentration (n=2 independent experiments). Error bars represent mean ± standard deviation; P values determined by t test or ANOVA with post-test multiple. ANOVA, analysis of variance; Only statistically significant comparisons (p<0.05) are highlighted.

Treatment of hepatocytes with a combination of oleic acid and palmitic acid (OA PA) in a 2:1 ratio induced steatosis in primary hepatocytes and cell lines. However, neither TLR5 inhibition nor addition of flagellin affected lipid accumulation (**Figure 3b, Supplementary Figure 4a, b**), expression of proteins associated with fatty acid oxidation (very long chain specific acyl-CoA dehydrogenase – ACADVL, medium-chain specific acyl-CoA dehydrogenase - ACADM and long-chain 3-hydroxyl-CoA dehydrogenase – HADHA, **Figure 3c,d,e, Supplementary Figure 4c-h**) or markers of ER stress (DNA damage-inducible transcript 3 (DDIT3) expression, **Figure 3f, Supplementary Figure 5a, b**)

OA PA-induced IL-8 production was TLR5-dependent in both primary and cell line hepatocyte cultures (**Figure 3a, Supplementary Figure 5c,d)** in which the absence of flagellin was confirmed (**Supplementary Figure 5e)**. TLR5 inhibition significantly reduced IL-8 production, while inhibition of TLR2 (pab hTLR2), TLR3 (CU-CPT 4a) and TLR4 (pab hTLR4) did not, in primary hepatocytes and both cell lines (**Figure 3a, Supplementary Figure 5c, d**). At lower concentrations of OA and PA, there was a dose-dependent synergistic effect with flagellin on IL-8 production (**Supplementary Figure 5f**), which was, again, reduced by TLR5 inhibition. Furthermore, OA PA did not induce IL-8 production in kidney (HEK) cells (**Supplementary Figure 5g**), suggesting a specific effect in hepatocytes. Together these results show that IL-8 is an injury-associated cytokine in human MASH and in vitro, TLR5 mediates both flagellin-induced and OA PA-induced IL-8 production.

### TLR5 mediates lipid-induced IL-8 production via the canonical MyD88 pathway

Treatment of the sensitive TLR5-linked NF-kB reporter assay system (HEK-Blue hTLR5) with OA PA resulted in no activation of the TLR5-NF-kB pathway (**Figure 4a**), suggesting that these FFAs are unlikely to be direct ligands for TLR5 and that any FFA-induced TLR5 ligand may be specific to hepatocyte injury. Furthermore, inhibition of lipid uptake into primary hepatocytes and HepG2 and Huh7 cell lines (**Figure 4b, Supplementary Figure 6a,b)** using a cocktail of inhibitors targeting key fatty acid transporters in hepatocytes (FATP2 ^26^ - lipofermata; FATP5 ^27^ - obeticholic acid, and CD36 ^28^ - SML2148) abolished IL-8 production in response to lipid loading, principally through FATP2 inhibition (**Figure 4c, Supplementary Figure 6c,d**). Consistent with current knowledge of TLR5 signalling to NF-kB activation and cytokine production, pharmacological inhibition of MyD88 and TAK1 but not TRIF significantly reduced IL-8 production in primary hepatocytes and both cell lines (**Figure 4d, e, Supplementary Figure 7a-d)**. Together, our data suggest that TLR5-mediated IL-8 production requires internalisation of FFAs into hepatocytes. Therefore, this implies the existence of secondary mediator(s) that activate TLR5 through its extracellular domain (TH1020 binds to the extracellular domain of TLR5) and via its known canonical pathway (**Figure 4f**).

**Figure 4.**
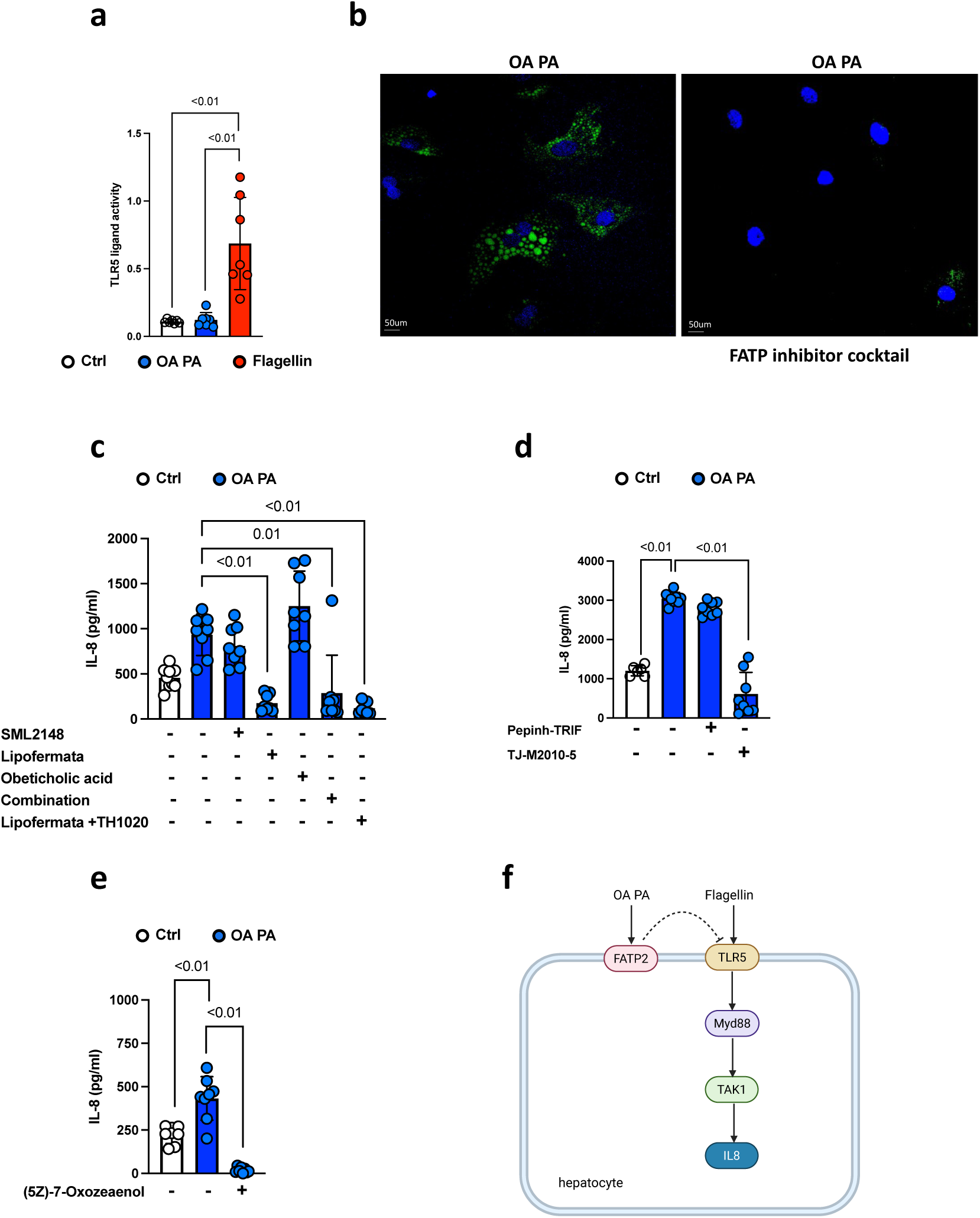
TLR5-mediated IL-8 production requires fatty acid internalisation into hepatocytes and signals via MyD88. **a**, HEK-Blue™ hTLR5 cells treated with oleic acid and palmitic acid (OA PA), flagellin for 24 hours. TLR5 ligand activity assessed by optical density measurements of SEAP production. n=3 independent experiments. **b**, Primary hepatocytes treated with OA PA for 24 hours in the presence or absence of fatty acid transporter protein inhibitor cocktail (10µM obeticholic acid (FATP5 inhibitor), 80µM SML2148 (CD36 inhibitor), 50µM Lipofermata (FATP2 inhibitor)). Representative images of lipid deposition quantified using BODIPY 493/503 staining (green), nuclei stained with DAPI (blue) via confocal microscopy (Scale bars 50µm). Experiments performed in triplicate for each condition. **c – e**, IL-8 concentrations in **c,** Huh7 cells treated with OA PA for 24 hours in the presence or absence of 16µM SML2148, 10µM Lipofermata, 2µM obeticholic acid, combination of 16µM SML2148, 10µM Lipofermata and 2µM obeticholic acid in a cocktail, 1µM TH1020 (TLR5 inhibitor) with 10µM Lipofermata (n=3 independent experiments). **d**, Primary hepatocytes treated with OA PA for 24 hours in the presence or absence of 40µM Pepinh-TRIF (TRIF inhibitor), 40µM TJ-M2010-5 (MyD88 inhibitor) (n=2 independent experiments) and **e**, HepG2 cells treated with OA PA for 24 hours in the presence or absence of 5µM (5Z)-7-Oxozeaenol (TAK1 inhibitor) (n=3 independent experiments). **f,** Schematic diagram depicting TLR5 signalling in hepatocytes (created using Biorender). Error bars represent mean ± standard deviation; P values determined by t test or ANOVA with post-test multiple comparisons. ANOVA, analysis of variance. Only statistically significant comparisons (p<0.05) are highlighted.

### TLR5 inhibition protects against lipid-induced HSC activation and collagen production in patient-derived human precision-cut liver slices (hPCLS)

Activated HSC predominantly produce collagen types I and III, which are the most abundant fibrillar collagens that accumulate during liver fibrosis ^29^ ^30^. Primary HSC isolated from a cohort of non-fibrotic non-steatotic human liver tissues (n=19 sampled from people undergoing hepatic resection surgery) did not express TLR5 by RNA sequencing (**Figure 5a**). Accordingly, neither flagellin nor OA PA directly induced TGFβ1 (marker of HSC activation) and pro-collagen 1α1 (precursor to collagen I) protein production in the LX2 HSC cell line (**Figure 5b, c**). However, conditioned media from HepG2 treated with OA PA did induce TGFβ1 and pro-collagen 1α1 production (**Figure 5d, e**). This induction was abrogated by inhibition of TLR5 in HepG2 before OA PA-mediated injury. This was confirmed using conditioned media from primary hepatocytes (**Supplementary Figure 7e**).

**Figure 5.**
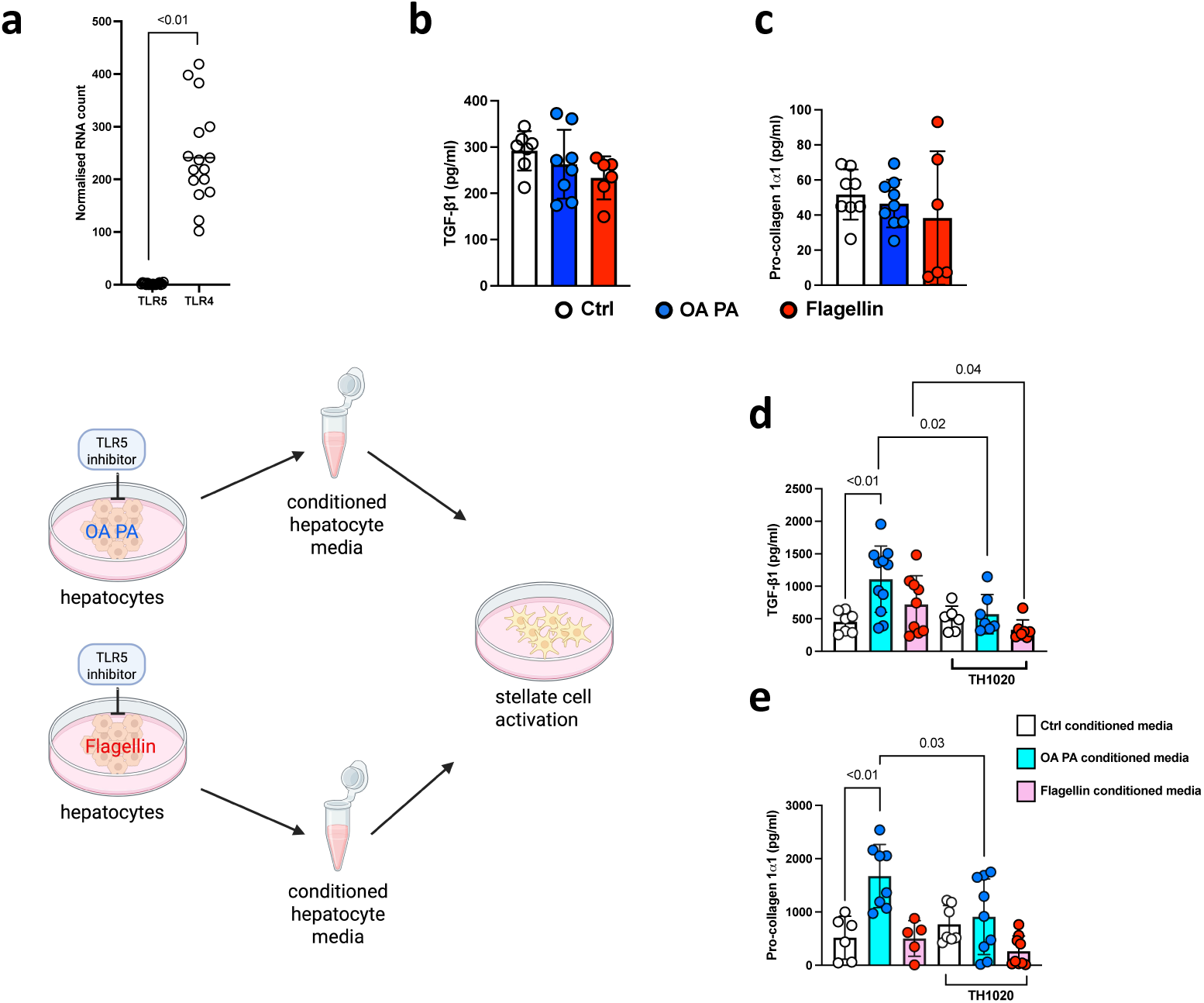
TLR5 inhibition on hepatocytes abrogates hepatic stellate cell activation a, Normalised RNA counts for TLR5 and TLR4 extracted from primary human hepatic stellate cells isolated from non-fibrotic, non-steatotic human livers (n=19). **b**, TGFβ1 concentration **c**, Pro-collagen 1⍺1 concentration, in LX2 cells treated with oleic acid and palmitic acid (OA PA), flagellin for 24 hours (n=3 independent experiments). **d**, TGFβ1 concentration, **e**, Pro-collagen 1⍺1 concentration, in LX2 cells treated with conditioned media from HepG2 cells treated with OA PA, flagellin in the presence or absence of 1µM TH1020 (TLR5 inhibitor) (n=3 independent experiments). Schematic diagram created using Biorender. Error bars represent mean ± standard deviation; P values determined by t test or ANOVA with post-test multiple comparisons. ANOVA, analysis of variance. Only statistically significant comparisons (p<0.05) are highlighted.

To further develop the human-relevance of these findings, the effect of TLR5 inhibition on OA PA-mediated injury was investigated in human precision-cut liver slices (hPCLS) taken from margins of hepatic resection surgery. hPCLS are uniform three-dimensional slices of fresh patient-derived liver tissue used as an ex vivo culture model. hPCLS retain the native architecture of liver tissue, extracellular matrix composition and cell-to-cell interactions making them more physiologically-relevant compared to 2D culture systems. Consistent with our findings in primary hepatocytes and cell lines, OA PA induced 2.8-fold increase in expression of collagen III compared to control (p<0.01). Pre-treatment of hPCLS with TLR5 inhibitor TH1020 resulted in 2.2-fold reduction in expression of collagen III (p<0.01, **Figure 6**). These findings support the human relevance of TLR5 in MASH-associated fibrosis, and indicate that it could inform future diagnostic approaches and, potentially, therapeutic targeting.

**Figure 6.**
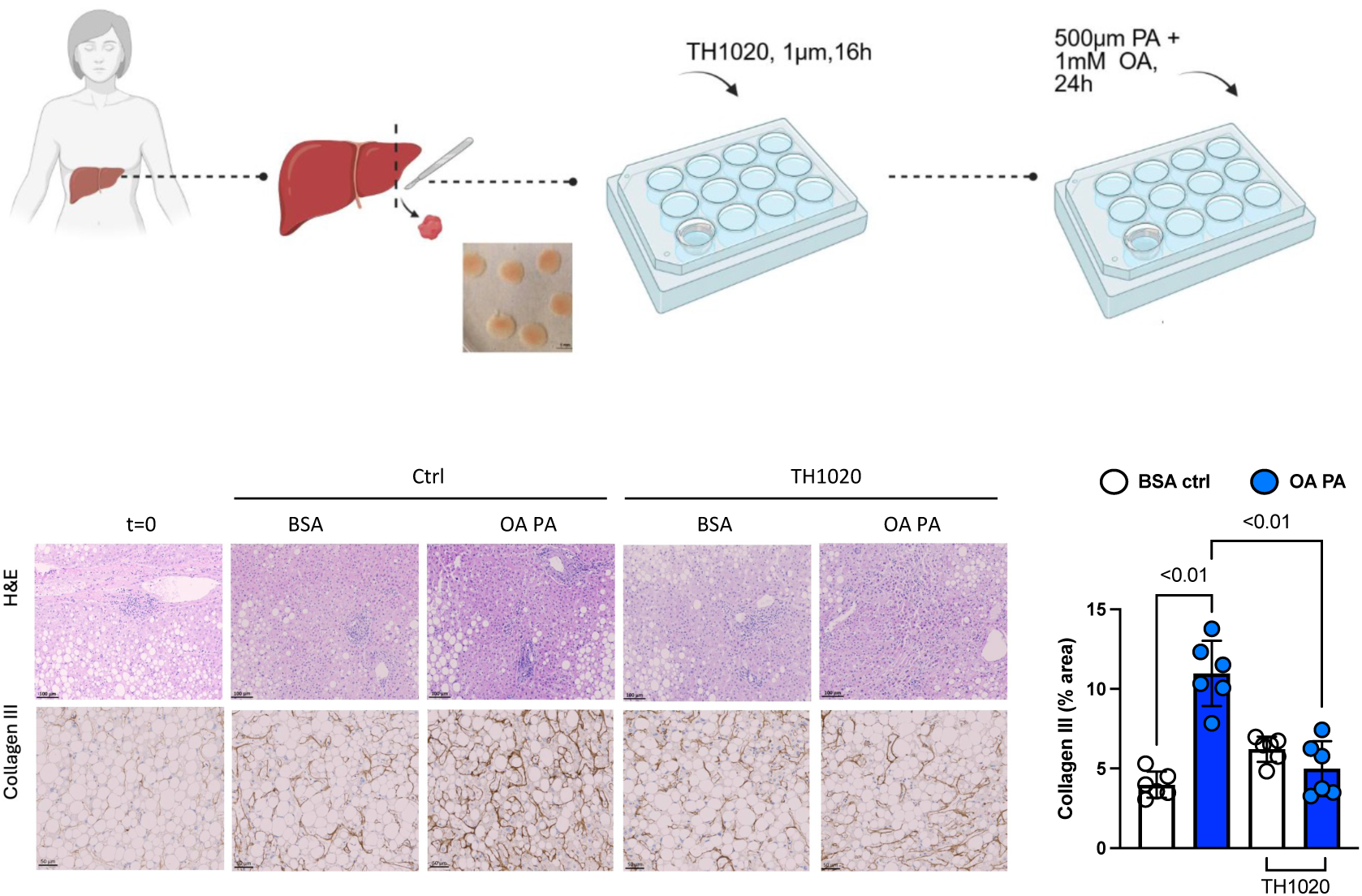
TLR5 inhibition reduces collagen deposition in human precision-cut liver slice model. Schematic representation of precision-cut liver slices (PCLS) setup, created in Biorender. PCLS were cultured for 16 hours in the presence or absence of 1µM TH1020 (TLR5 inhibitor) and treated with bovine serum albumin (BSA) or oleic acid and palmitic acid (OA PA) for 24 hours (n=3). Representative immunohistochemistry images H&E and Collagen III quantification (% area) of PCLS shown. Scale bar 100µM and 50µM. Error bars represent mean ± standard deviation; P values determined by t test or ANOVA with post-test multiple comparisons. ANOVA, analysis of variance. Only statistically significant comparisons (p<0.05) are highlighted.

## Discussion

Hepatic Toll-like receptors (TLRs) are sensors of pathogen- and damage-associated molecular patterns and are crucial gatekeepers of liver health. TLR5 expression is elevated in multiple independent MASLD cohorts with advanced MASH fibrosis and associated with adverse patient outcomes. Moreover, the combination of immunofluorescence, spatial transcriptomics and single cell RNAseq together indicate that elevated hepatic TLR5 expression is principally within hepatocytes. TLR5’s cognate ligand flagellin, is elevated in plasma from individuals with advanced MASH fibrosis and reduces with liver disease improvement following metabolic surgery. Mechanistically, hepatocyte TLR5 activation plays a central role in regulating pro-inflammatory cytokine production (IL-8) and fibrogenesis through indirect HSC activation. In the absence of flagellin, fatty acid transporter 2 (FATP2)-mediated internalisation of OA and PA leads to indirect activation of TLR5 likely through its extracellular domain. This uncovers a novel signalling interaction between TLR5 and fatty acids mediated through FATP2, which is highly expressed in the liver to facilitate uptake of long chain fatty acids ^31^ ^32^. Whether fatty acids themselves also act as ligands for TLRs remains unclear ^33^ ^34^ ^35^ ^36^ and work is ongoing to identify ligand-TLR5 receptor interactions in human MASH. We propose that inhibition of the TLR5 pathway has two parallel mechanisms that can mitigate hepatocyte injury: one elicited by PAMP-driven (flagellin) stimulation and the other (indirectly) by lipid injury, which in turn suppresses subsequent fibrogenesis mediated by HSC activation.

Flagellin is a virulence factor that confers bacterial mobility within the intestine facilitating access to niches conducive to colonisation, invasion and host inflammatory responses ^37^. In the liver, elevated flagellin levels have been associated with cell injury ^38^. Increased stool and plasma flagellin concentrations are observed in individuals with obesity ^39^ and type 2 diabetes^40^. Emerging evidence suggests that flagellin-mediated activation of TLR5 plays a critical role in pancreatic β-cell inflammation and dysfunction ^40^. Further work is needed to decipher the association between metabolic syndrome and enrichment of flagellated bacteria in the gut. Structurally, flagellins vary across different bacterial species and differ in TLR5 binding capacity ^41^ ^42^. Which specific flagellated bacteria/flagellins activate hepatic TLR5 signalling in MASH fibrosis is currently unclear. In this study, we used commercially available purified flagellin to avoid contamination from other bacterial components such as LPS to activate TLR5 signalling in vitro. Future studies using isolated flagellated bacteria/flagellin from liver tissue are warranted.

Clinically, hepatic TLR5 expression or plasma flagellin may be useful biomarkers for advanced MASH fibrosis and disease regression and therefore could be used to assess treatment responses to novel emerging pharmacotherapies in MASH. Further work is needed to determine their clinical utility in these settings in larger independent patient cohorts. It is reasonable to hypothesise that TLR5 inhibition is an actionable therapeutic strategy, although TLR5 is ubiquitously expressed ^43^ and its inhibition may lead to unintended adverse effects within the liver or in extra-hepatic tissues. Inhibition of downstream kinases (such as TAK1) or modulation of the gut microbiome to reduce flagellin or flagellated bacterial species may offer alternative strategies to modulate TLR5 signalling.

We acknowledge limitations of the current study. Whilst the combination of OA and PA is a widely-accepted model of lipid injury in hepatocytes ^44^ ^45^, the effect of other fatty acids or lipid species in mediating TLR5 signalling was not investigated. Longitudinal reduction in plasma flagellin may be specific to metabolic surgery itself due to changes in microbiome composition following structural alterations to the proximal gastrointestinal tract or reversal of metabolic syndrome rather than improvement in liver injury alone. The TH1020 inhibitor that was used in the current study targets the extracellular domain of TLR5 and further insight into binding patterns and mechanisms may be gained by wider pharmacologically-focussed experiments.

In conclusion, we show that activation of hepatic TLR5 signalling is directly associated with adverse patient outcomes in human MASH. Mechanistically, inhibition of TLR5 reduces lipid and flagellin-induced hepatocyte injury via MyD88-TAK1-IL-8 pathway, and abrogates HSC activation. Thus, pharmacologically targeting TLR5 signalling in hepatocytes may offer a novel therapeutic approach for MASH-associated fibrosis.

## Methods

### MASLD participants

Apart from re-analysis of published studies, this study included participants over the age of 18 attending the hepatology outpatient clinics at Barts Health NHS Trust or for elective metabolic surgery at Homerton University Hospital Foundation Trust (**Supplementary Table 1** for further participant details and samples used in this study). Participants provided written informed consent at enrolment. The studies were approved by local ethics Committee (reference numbers 18/LO/1759, 14/WA/1142) and performed in compliance with the Declaration of Helsinki. Participants ranging from healthy controls to MASH cirrhosis based on imaging (ultrasound, CT, MRI, elastography) and/or liver histology obtained either per-protocol at the time of metabolic surgery or in individuals with suspicion of MASH fibrosis based on elevated non-invasive liver fibrosis markers. All liver biopsies were reported by a single experienced histopathologist and were summarised according to the National Institutes of Health NASH clinical research network (Kleiner) criteria ^46^. Participants with evidence of other non-MASLD liver diseases, alcohol consumption > 14 units per week (women) and 21 units per week (men), taking concomitant immunosuppressive medications, non-type-2 diabetes or history of previous liver decompensation events were excluded.

Liver biopsy tissue used in this study was derived intra-operatively from participants undergoing elective metabolic surgery and obtained using Max-CoreTM Disposable Core Biopsy Instrument 18G x 20cm though laparoscopic port directly over the liver under direct vision by the operating surgeon and stored in RNAlater solution at −80°C until RNA extraction. Peripheral blood was collected on the day of study consent or on the day of metabolic surgery in sodium heparin blood collection tubes and centrifuged at 2500 revolutions per minute for 10 minutes to extract plasma which was immediately frozen at −80°C. Repeat venesection was performed in participants undergoing metabolic surgery following weight loss and repeat plasma extraction was performed. Stool samples were collected on the day of study consent from participants and immediately snap frozen in cryovials at −80°C.

### Independent international MASLD cohorts

Reported TLR5, TLR4 and TLR2 gene expression from whole liver tissue was compared according to liver disease severity in three independent biopsy-proven MASLD cohorts. Cohort 1: SteatoSITE cohort, n = 408 participants (control, n = 31; steatosis, n = 49; MASH F1-2, n = 148; MASH F3-4, n = 180). Cohort 2: University of Cambridge (UCAM), Virginia Commonwealth University (VCU) combined cohort, n = 136 participants (control, n = 4 and MASLD, n = 132). (mild (F0), n = 52; moderate (F1–F2), n = 50; severe (F3–F4), n = 30). Cohort 3: Newcastle University (EPoS) consisted of 168 MASLD participants ((mild (F0), n = 47; moderate (F1–2), n = 64; severe (F3–4), n = 57). Data from cohort 1 was extracted by SteatoSITE team from data published by Kendall et al ^47^. Data from cohort 2 and 3 were extracted from publicly available dataset by Vacca et al ^22^.

### Treatment naïve non-obese chronic hepatitis B participants

Plasma samples were obtained from participants over the age of 18 attending the hepatology outpatient clinics at Barts Health NHS Trust with confirmed treatment naïve chronic hepatitis B infection. Participants provided written informed consent at enrolment. The studies were approved by local ethics Committee (reference numbers 17/LO/0266) and performed in compliance with the Declaration of Helsinki. All participants had body mass index measurement and non-invasive fibrosis assessment by transient elastography within 12 months of study recruitment.

### Patient-related outcomes analysis (SteatoSITE)

To determine effects on survival and liver-related events in the SteatoSITE MASLD cohort, normalised counts per minute of TLR5 from the biopsy subset of cases in the SteatoSITE data commons were used in survival analysis using R (v.4.3.0) in RStudio (v.2023.12.0 build 369) and the ‘survminer’ package (v.0.4.9). The optimal cutpoint for normalised TLR5 counts was separately calculated for overall survival, and hepatic decompensation (when a first coding of any component of the composite outcome occurred after the biopsy date and with death as a competing risk) using surv_cutpoint, applying maximally selected rank statistics of the ‘maxstat’ package (v.0.7-25) with a minimum proportion of 0.25. Kaplan–Meier estimator curves of all-cause mortality were compared by regular log-rank testing with weights = 1.

### Shotgun metagenomics and stool flagellin gene load analysis

Patient stool samples stored in cryovials at −80°C underwent shotgun metagenomics sequencing. The protocol involved DNA extraction using Qiagen PowerFecal DNA kit, DNA sequencing library preparation and then sequencing using the Illumina HiSeq4000 platform. A pseudo custom reference genome specific to flagellin gene (FliC) was created using NCBI gene browser across multiple bacterial species (https://www.ncbi.nlm.nih.gov/genome/?term=FliC. This was used to assess the number of reads originating from *FliC* gene sequences from multiple bacterial species. Alignment was performed using BWA (v0.7.17) and verified using Bowtie2 (v2.4.5). Reads were quantified using Samtools (v1.10) flagstat (https://www.htslib.org/doc/samtools-flagstat.html).

### Cell culture and treatments

Cells used in this study for in vitro experiments include primary hepatocytes from healthy donors (BioIVT Human Cryoplateable Hepatocytes, ref: MEF), HepG2, Huh7 and LX2 cell lines. Primary hepatocytes were cultured in INVITROGRO™ CP Medium supplemented with TORPEDO™ Antibiotic Mix according to the manufacturer’s protocol. HepG2 and Huh7 cells were cultured in DMEM media (Gibco) supplemented with 10% heat-inactivated fetal bovine serum (Gibco), L-glutamine, 100 Units penicillin and 100 μg streptomycin per mL. LX2 cells were cultured in DMEM media (Gibco) supplemented with 2% heat-inactivated fetal calf serum (Gibco), L-glutamine, 100 Units penicillin and 100 μg streptomycin per mL. Experiments were conducted in 24-well plates at 100,000 cells per mL of DMEM media and incubated at 37°C. Confluent hepatocytes were treated with fatty acid–free BSA (MilliporeSigma), 1% BSA-conjugated oleic acid (OA 1000μM), palmitic acid (PA 500μM) at 2:1 ratio (OA PA), flagellin (FLA-ST Ultrapure, 100ng/mL, Invivogen) or combination (OA PA + flagellin) for 24 hours. We used the TLR5 inhibitor TH1020 (1μM, Sigma-Aldrich). Seeded cells were treated overnight (18 hours) before the cell media was changed for in vitro experiments. See **Supplementary Table 4** for further details of other inhibitors used in this study.

HEKBlue-hTLR5 reporter cell and HEKBlue-hTLR null reporter cell lines (Invivogen) were grown in DMEM supplemented with 10% FCS, 2 mM L-glutamine, 100 µg/mL Normocin in the presence of selection antibiotics. Stimulation with a TLR ligand activates NF-κB and AP-1-induced production of secreted embryonic alkaline phosphatase (SEAP), which was directly measured directly using a spectrophotometer at 620-655 nm, and then be subtracted from background SEAP produced from the parental cell line HEK293 Null cells.

### Measurement of cytokines

Cytokine concentrations were determined by ELISA (enzyme-linked immunosorbent assay) according to manufacturer instructions. The ELISAs kits used in this study are listed in **Supplementary Table 5.** Plasma IL-8 quantification from MASLD participants were determined Luminex assay (R&D, Human Magnetic Luminex Assay) according to manufacturer instructions.

### Immunofluorescence Staining

Liver biopsy sections mounted on glass slides were de-paraffinized using Histoclear II solution (National Diagnostics) followed by re-hydration using a series of Ethanol concentrations (100%, 90% and 70%) before being washed with water. Sections were heated to 95°C for 3 rounds of 3 minutes in antigen retrieval solution (ab64236, Abcam) and allowed to cool to room temperature. Sections were initially blocked in 3% BSA diluted in PBS +0.1% Tween 20 (PBS-T; P1379-100ML, Sigma) for 1 hour at room temperature. The sections were incubated with primary antibodies (**Supplementary Table 6**) overnight (diluted in 1% BSA in PBS-T). They were then washed 3 times with PBS-T and incubated with appropriate secondary antibodies (**Supplementary Table 6**) for 1 hour at room temperature. The sections were washed and mounted with FluoroMount G (00-4958-02, Invitrogen). Images were acquired using an inverted Nikon Spinning) disc SoRa microscope. Images were analysed with ImageJ (NIH) and Volocity (Version 6.3, PerkinElmer).

### BODIPY 493/503 staining

Hepatocytes were fixed using 4% PFA for 30 minutes at room temperature, followed by three PBS washes. Hepatocytes were then permeabilised with 0.1% Triton X-100 and incubated with 2 μM BODIPY 493/503 staining solution in PBS for 15 minutes at RT followed by Hoechst 33342 (1μg/ml) or DAPI (1μg/ml) and further incubation (1 hour at room temperature), then imaged under confocal microscope (Zeiss 880 Laser Scanning Confocal Microscope with Fast Airyscan and Multiphoton (inverted). Stained cells were analysed using ImageJ software to quantify relative intensity/Hoechst or DAPI.

### Quantitative Polymerase Chain Reaction Assay

Cells were lysed with TRIzol Reagent (Thermo Fisher) and total RNA was isolated following the manufacturer’s instructions. RNA concentrations were measured spectrophotometrically using NanoDrop 2000 (Thermo Fisher) and 3 μg was used for each cDNA preparation. cDNA was reverse transcribed using the Superscript IV protocol (Invitrogen) The RT conditions for each cDNA amplification were 42 °C for 15 minutes, 85 °C for 5 seconds, and the cDNAs amplified were stored at −20 °C. Gene expression analysis was performed by qPCR using the rotor-gene SYBR green PCR kit (Qiagen) according to the manufacturer’s protocol. The relative expression levels of the studied genes were standardised to the GAPDH as control housekeeping gene as an internal control in the same sample and stated as the fold difference in expression using the 2-ΔΔCt method.

### RNA sequencing of primary human hepatic stellate cells

Primary human hepatic stellate cells (hHSCs) were isolated from non-fibrotic, non-steatotic human livers as previously described by Rombouts et al ^48^. Study was approved by National Ethics Service Research Ethics Service (NRES), Research Ethics Committee (REC) number: 21/WA/0388. RNA was extracted from liquid-nitrogen cryopreserved hHSCs using the RNeasy Plus Mini kit (Qiagen) according to the manufacturer’s instructions. Quantification and purity assessment were performed on a Nanodrop 2000 (ThermoFisher). RNA sequencing was performed in collaboration with UCL Genomics. Sample quality was further assessed on Agilent TapeStation. KAPAmRNA HyperPrep kit was used for library preparation and samples were sequenced using paired-end reads with unique dual indexes on the NovaSeq instrument (Illumuna, San Diego, US). Data normalisation and quality control were performed by UCL Genomics.

### Precision cut liver slice model

Human liver samples were procured from areas located at least 10 cm away from both the tumour site and the surgical margin to ensure the collection of non-malignant tissue. The study included one patient with steatohepatitis at baseline. Ethical approval was granted by the UZ/KU Leuven Ethics Committee (S67418). From the resection specimen, three spatially distinct regions of liver tissue were isolated. Histological assessment of the specimens was confirmed by an experienced liver pathologist. Samples were sectioned into 250 μm slices and cultured on transwell inserts with 8 μm pores, placed in 12-well plates (Greiner Bio-One). Liver slices were maintained at 37 °C with continuous agitation in Williams Medium E, supplemented with 1% penicillin–streptomycin and L-glutamine (Sigma-Aldrich). For TLR5 inhibition, liver slices were incubated with 1 µM TLR5 antagonist, TH1020, (HY-116961-5, MedChemexpress) for 16 hours. Lipid loading was performed using either a 1% solution of fatty acid–depleted bovine serum albumin (BSA; Sigma-Aldrich) as vehicle control or a BSA-conjugated mixture of 500 µM palmitic acid and 1000 µM oleic acid for 24 hours. Following treatment, tissues were fixed, paraffin-embedded, and processed as FFPE blocks for immunostaining.

### Immunohistochemistry staining from PCLS

FFPE tissue sections were immunostained with rabbit polyclonal antibody directed against human Collagen Type III (rabbit IgG, ref: 22734-1-A, Proteintech, 1/2000) using the fully automated BOND Max system from Leica Microsystems and a BOND Polymer Refine Detection kit (#DS9800). Slides were scanned with the AxioScan Z1 slide scanner (Zeiss). For collagen III quantification, six representative regions of interest (ROIs) were selected per condition, and quantitative analysis was performed using ImageJ software (version 1.54p).

### CosMx™ Spatial Molecular Imaging

CosMx™ spatial molecular imaging (Bruker Spatial Biology) was performed using the CosMxTM Human Universal Cell Characterization RNA Panel (1000-plex) supplemented with 14 custom genes of interest on n=8 biopsy samples and n=2 explant liver samples from individuals with end-stage MASH at the University Hospitals Leuven. Ethical approval was granted by the UZ/KU Leuven Ethics Committee (S67418). Morphology was visualised using B2M/CD298, PanCK, CD45 and CD68, combined with DAPI.

Samples were prepared as follows: 5 µm-thick FFPE sections were mounted on VWR Superfrost Plus Micro slides (cat# 48311-703) were baked for 4 hours at 37°C and stored at 4°C, then processed 7 days later using the Leica Bond RXm system (Leica) for staining and probe incubation according to CosMx guidelines. RNA target readout was performed on the CosMx SMI instrument as described previously (https://pubmed.ncbi.nlm.nih.gov/36203011/). Cloud-based AtoMx™ Spatial Informatics Platform (Bruker Spatial Biology) was used to export data and images for Seurat (v5.1.0). (https://pubmed.ncbi.nlm.nih.gov/37231261/). The data was merged, SCT normalised, followed by PCA and UMAP calculation using 30 dimensions with Seurat default parameters for CosMx data. Louvain clustering was performed with a resolution of 0.3, and clusters were combined and labelled based on canonical markers. The hepatocyte cluster (230,359 cells) was subsetted from the data using the subset function, then normalised and scaled, followed by PCA and UMAP calculation using 20 dimensions with Seurat default parameters. Seurat v5 was used for visualisation with default functions, and areas of interest captured with the Crop function.

### Single-nucleus RNA-sequencing (snRNA-seq)

Previously analysed publicly available snRNA-seq data (GSE202379) ^23^ was accessed through https://www.mohorianulab.org/shiny/vallier/LiverPlasticity_GribbenGalanakis2024/. Gene coexpression analysis was used to identify total number of single cells expressing TLR5 and ALB (albumin, hepatocyte marker).

### Statistical analysis

All data are presented as mean ± standard deviation. Statistical significance was determined using t-test or ANOVA with post-test multiple comparisons using GraphPad Prism 10.0.2. In all cases, A P-value < 0.05 was considered statistically significant and only statistically significant comparisons were highlighted in figures.

---

**Supplementary Figure 1.**
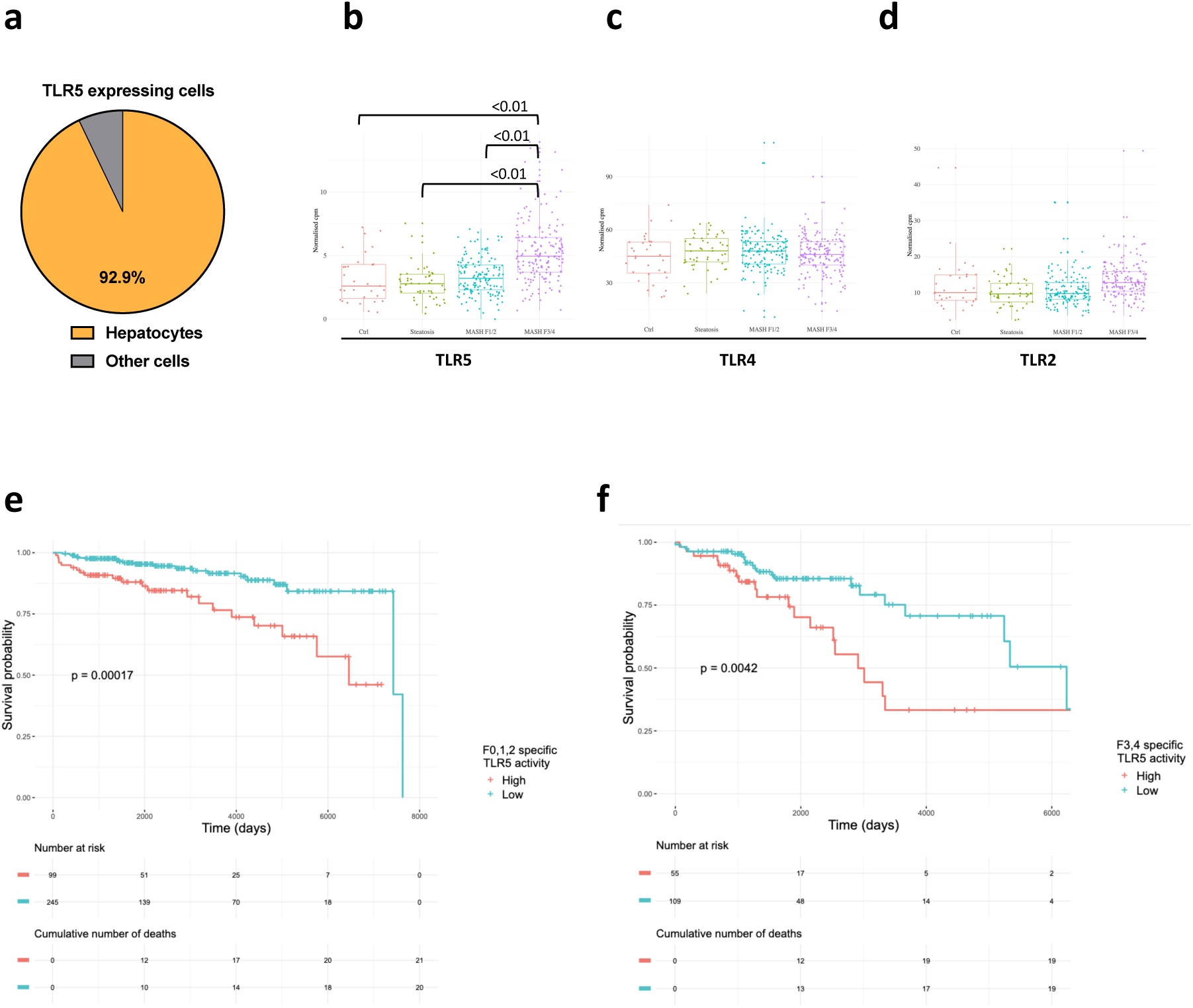
**a**, Publicly-available single nucleus RNA sequencing data from 47 liver biopsies from individuals with different stages of MASLD (Gribben et al. Nature 2024, PMID 38778114 ^23^) were analysed. Pie chart demonstrates proportion of TLR5 expressing single cells (n=1609) that co-expressed albumin (ALB, n=1495), equating to 92.9% derived from albumin-expressing hepatocytes. **b**, Liver TLR5 mRNA expression in SteatoSITE cohort divided according to histological MASLD disease severity. **c**, Liver TLR4 mRNA expression in SteatoSITE cohort divided according to histological MASLD disease severity. **d,** Liver TLR2 mRNA expression in SteatoSITE cohort divided according to histological MASLD disease severity. **e**, All-cause mortality by TLR5 expression in SteatoSITE cohort with MASH fibrosis stage 0 – 2. **f**, All-cause mortality by TLR5 expression in SteatoSITE cohort with MASH fibrosis stage 3 – 4. Kaplan–Meier curves for all-cause mortality stratified by hepatic TLR5 expression. P values determined by t test or ANOVA with post-test multiple comparisons. ANOVA, analysis of variance. Only statistically significant comparisons (p<0.05) are highlighted.

**Supplementary Figure 2.**
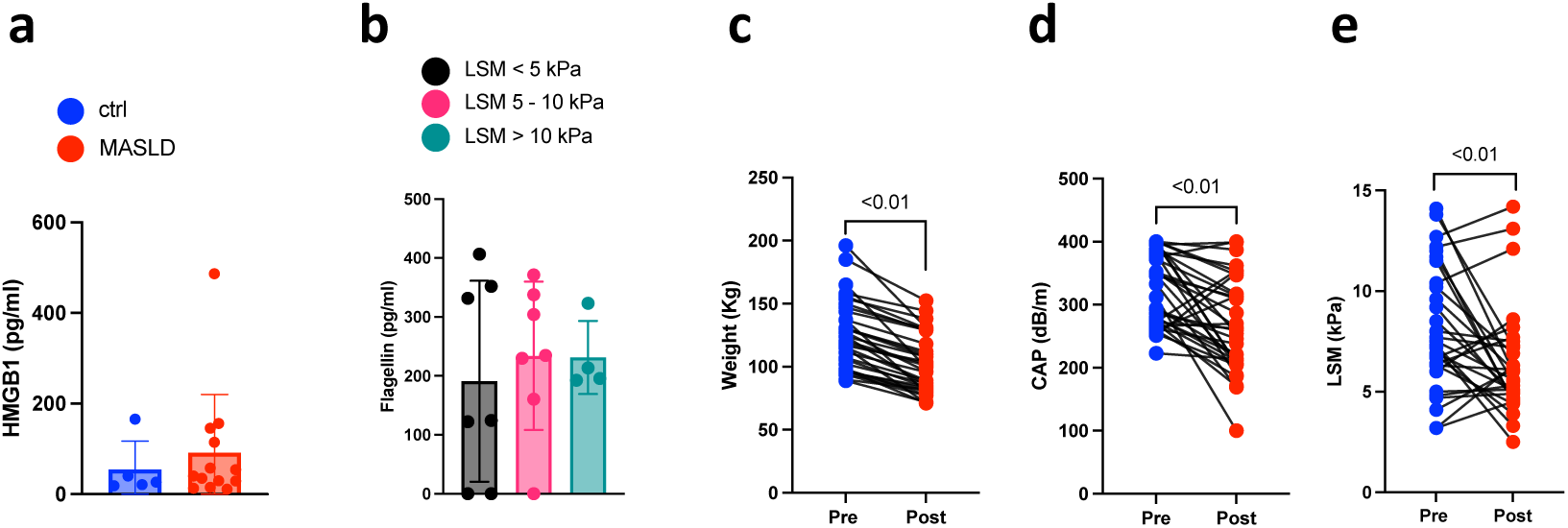
**a**, Plasma HMGB1 concentration in 18 participants (control n=5, MASLD n=13). **b**, Plasma flagellin concentration in 18 non-obese individuals with treatment naïve chronic hepatitis B infection, divided according to liver stiffness measurement (LSM) (LSM <5kPa n=7, LSM 5-10 kPa n=7, LSM >10kPa n=4). **c,d,e,** Longitudinal change following metabolic surgery in MASLD participants in **c**, Weight (kg); **d**, Controlled attenuation parameter (CAP), **e**, Liver stiffness measurement (LSM). P values determined by t test or ANOVA with post-test multiple comparisons. ANOVA, analysis of variance. Only statistically significant comparisons (p<0.05) are highlighted.

**Supplementary Figure 3.**
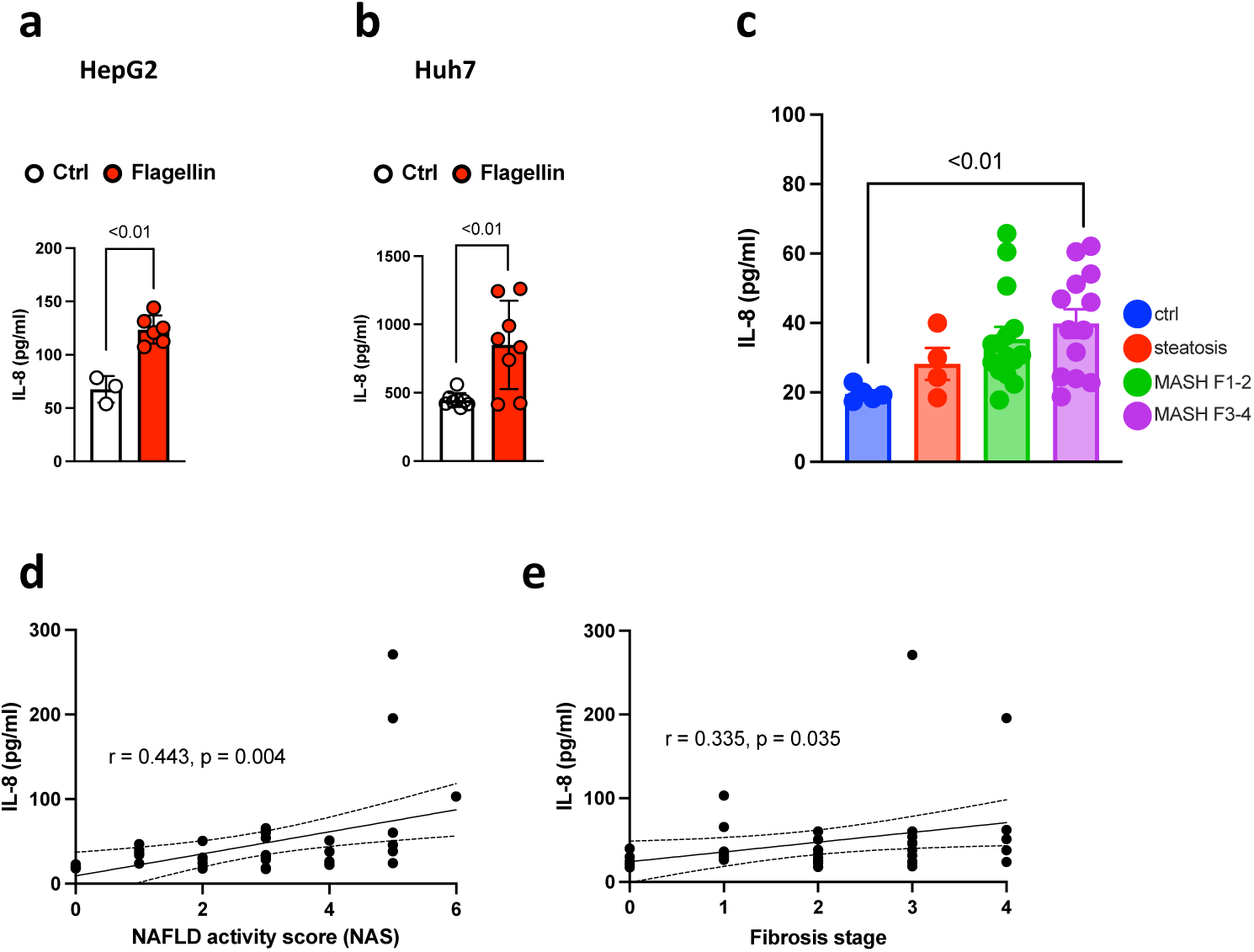
IL-8 concentration in **a,** HepG2 cells; **b**, Huh7 cells, following treatment with flagellin for 24 hours (n=3 independent experiments). **c**, Plasma IL-8 concentration from 37 participants by histological stage (control n=5, steatosis n=4, MASH F1-2 n=15, MASH F3-4 n=13). **d**, Pearson correlation of plasma IL-8 concentration and NAFLD activity score (NAS); **e**, Pearson correlation of plasma IL-8 concentration and histological fibrosis stage. Error bars represent mean ± standard deviation; P values determined by t test or ANOVA with post-test multiple comparisons or Pearson’s correlation coefficient. ANOVA, analysis of variance. Only statistically significant comparisons (p<0.05) are highlighted.

**Supplementary Figure 4.**
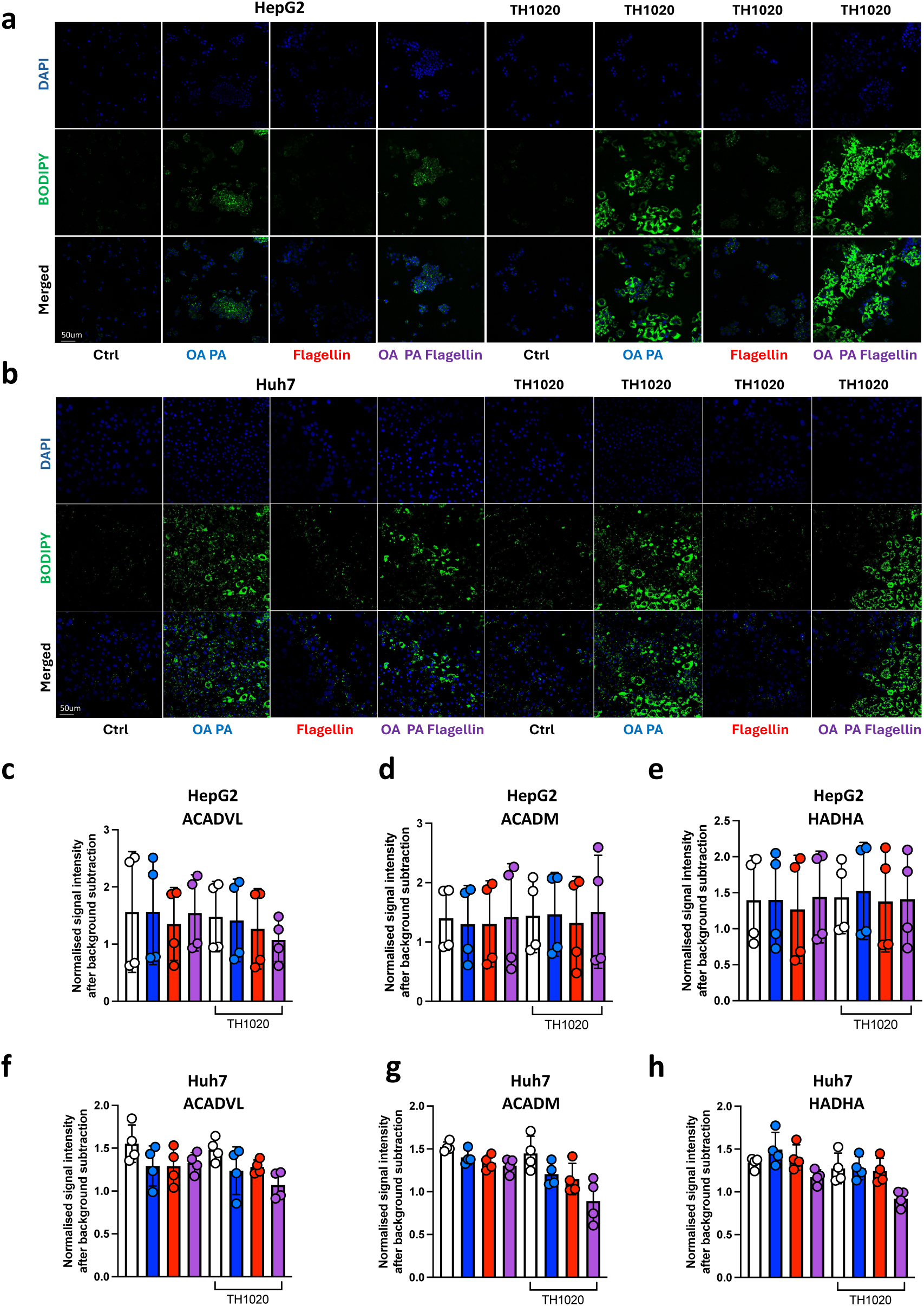
**a**, HepG2 cells, **b**, Huh7 cells, treated with oleic acid and palmitic acid (OA PA), flagellin or combination (OA PA + flagellin) for 24 hours in the presence or absence of 1µM TH1020 (TLR5 inhibitor). Representative images of lipid deposition quantified using BODIPY 493/503 staining (green), nuclei stained with DAPI (blue) via confocal microscopy (Scale bars 50µm). Experiments performed in triplicates for each condition. **c – h,** Expression of fatty acid oxidation enzymes (**c – e** in HepG2 cells, **f – h** in Huh7 cells) of very long chain specific acyl-CoA dehydrogenase – ACADVL, medium-chain specific acyl-CoA dehydrogenase - ACADM and long-chain 3-hydroxyl-CoA dehydrogenase – HADHA following treatment with OA PA, flagellin or combination (OA PA flagellin) for 24 hours in the presence or absence of 1µM TH1020 (n=2 independent experiments). Error bars represent mean ± standard deviation; P values determined by t test or ANOVA with post-test multiple comparisons. ANOVA, analysis of variance. Only statistically significant comparisons (p<0.05) are highlighted.

**Supplementary Figure 5.**
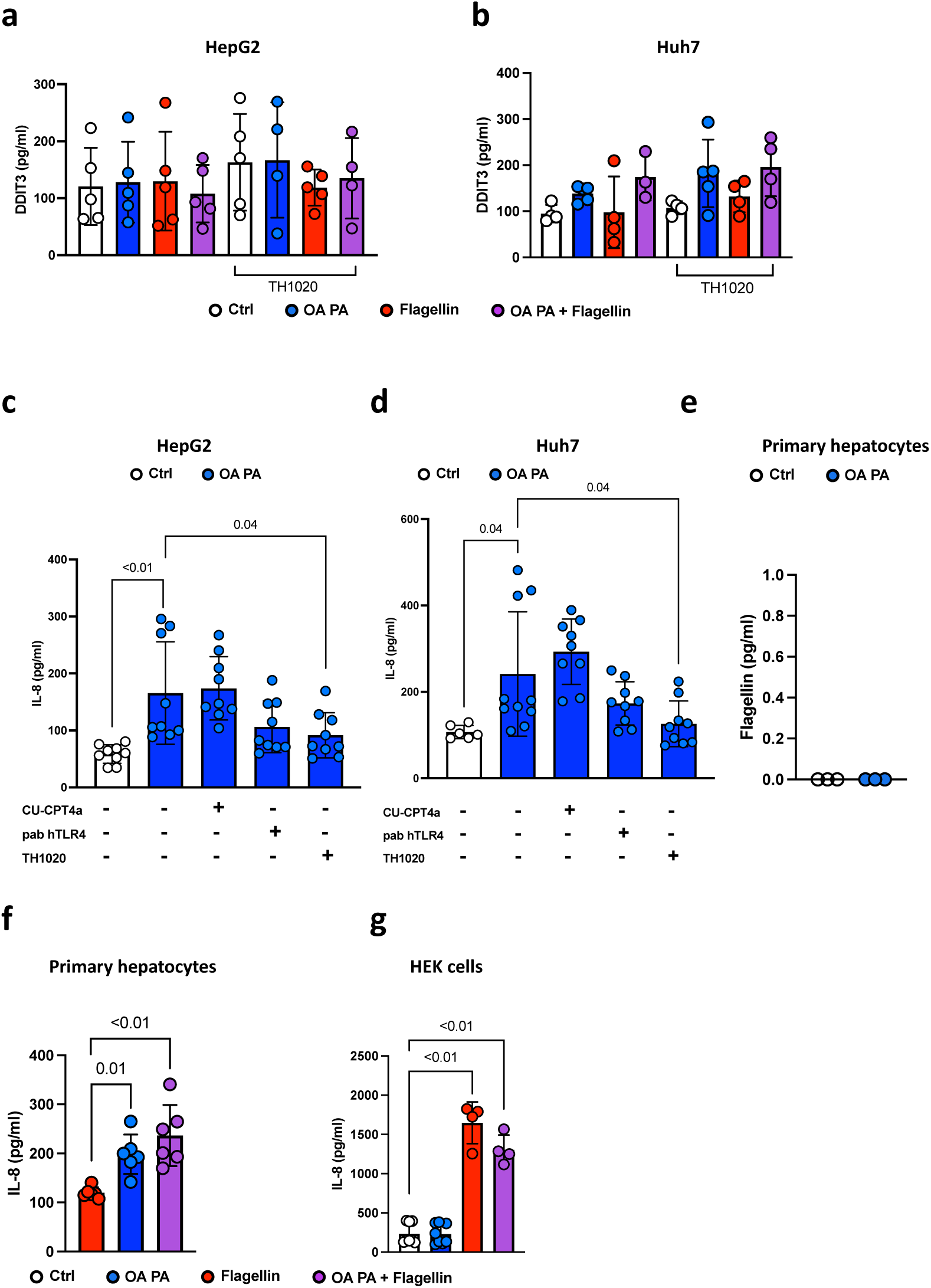
DNA damage-inducible transcript 3 (DDIT3) concentration in **a**, HepG2 cells; **b**, Huh7 cells, treated with oleic acid and palmitic acid (OA PA), flagellin or combination (OA PA + flagellin) for 24 hours in the presence or absence of 1µM TH1020 (TLR5 inhibitor) (n=2 independent experiments). **c**,**d**, IL-8 concentration in **c**, HepG2 cells, **d**, Huh7 cells, treated with OA PA for 24 hours in the presence or absence of 27µM CU-CPT4a (TLR3 inhibitor), 5µg/mL PAb hTLR4 (TLR5 inhibitor), 1µM TH1020 (n=3 independent experiments). **e**, Flagellin concentration conditioned media from primary hepatocytes treated for 24 hours treatment with OA PA compared to control (n=3 independent experiments). **f**, IL-8 concentration in primary hepatocytes treated with 250µM oleic acid 125µM palmitic acid (OA PA), 25ng/mL flagellin or combination (OA PA + flagellin) for 24 hours (n=2 independent experiments). **g**, IL-8 concentration in HEK cells treated with OA PA, flagellin, and combination (OA PA + flagellin) for 24 hours (n=3 independent experiments). Error bars represent mean ± standard deviation; P values determined by t test or ANOVA with post-test multiple comparisons. ANOVA, analysis of variance. Only statistically significant comparisons (p<0.05) are highlighted.

**Supplementary Figure 6.**
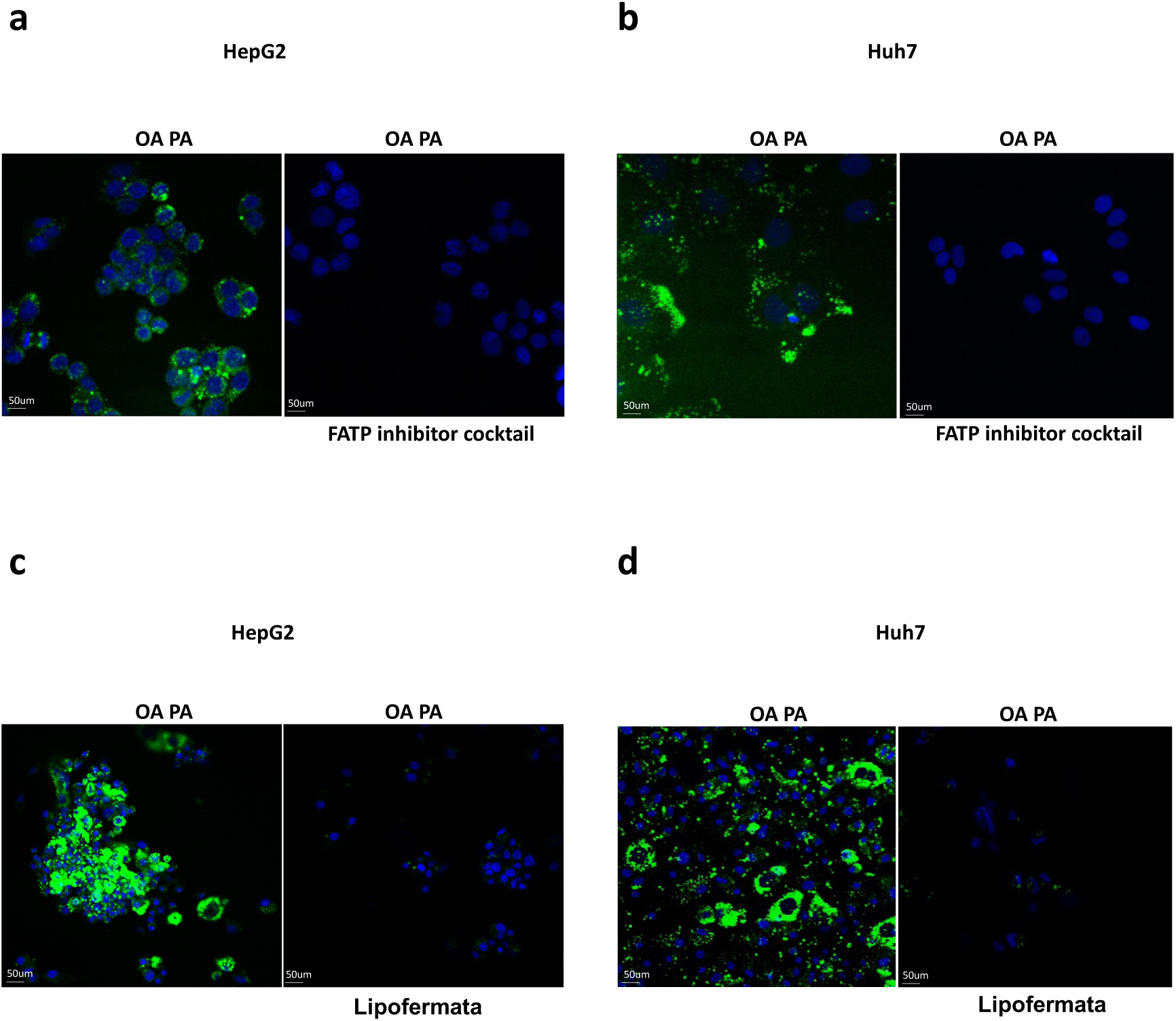
**a**-**d**, Representative images of lipid deposition quantified using BODIPY 493/503 staining (green), nuclei stained with DAPI (blue) via confocal microscopy (Scale bars 50µm) in **a,** HepG2 cells; **b**, Huh7 cells, treated with oleic acid and palmitic acid (OA PA) for 24 hours in the presence or absence of fatty acid transporter protein inhibitor cocktail (2µM obeticholic acid (FATP5 inhibitor), 16µM SML2148 (CD36 inhibitor), 10µM Lipofermata (FATP2 inhibitor)). **c**, HepG2 cells, **d**, Huh7 cells, treated with OA PA for 24 hours in the presence or absence of 10µM Lipofermata. Experiments performed in triplicates for each condition.

**Supplementary Figure 7.**
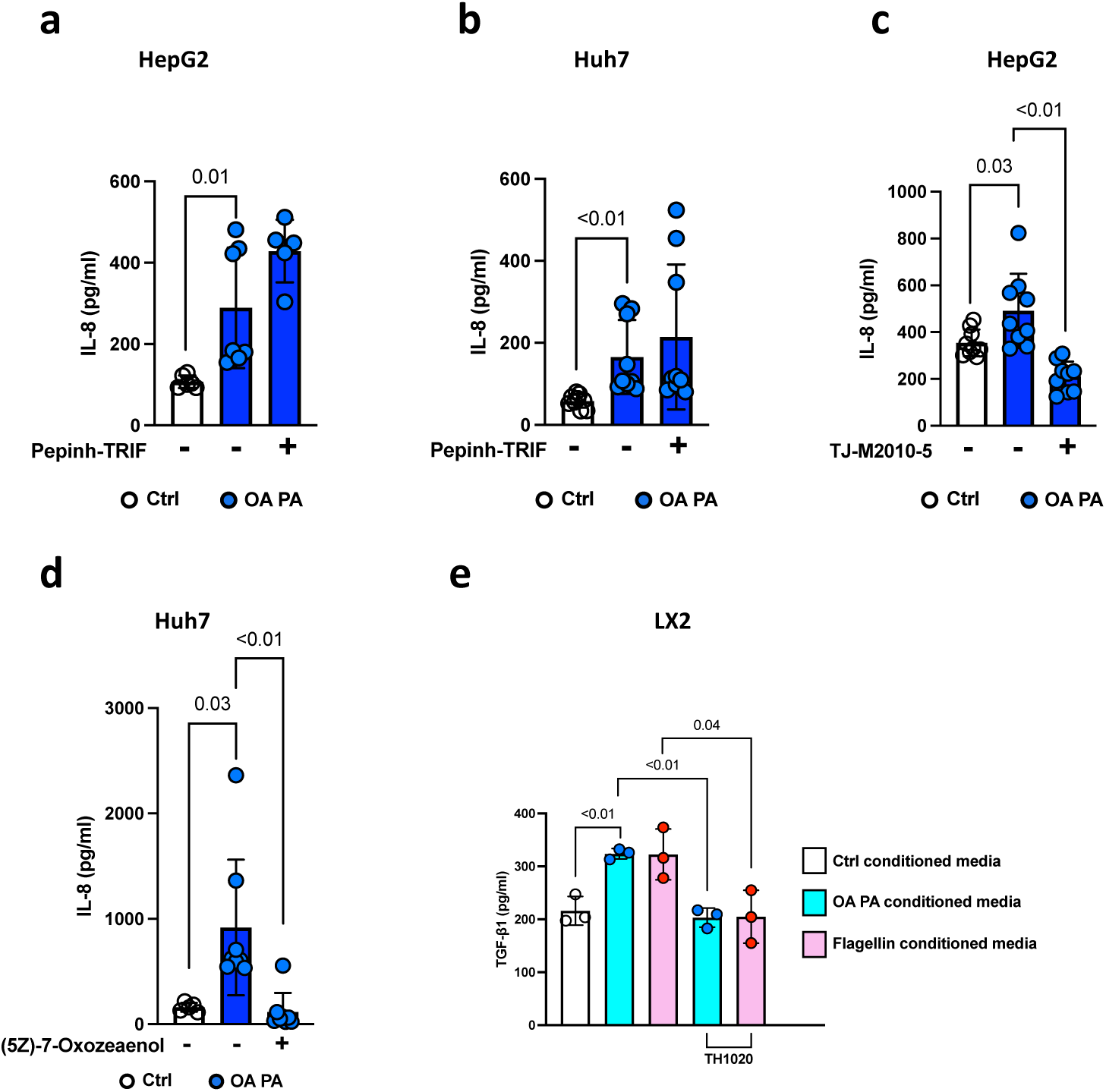
**a**,**b**, IL-8 concentration in **a**, HepG2 cells; **b**, Huh7 cells, treated with oleic acid and palmitic acid (OA PA) for 24 hours in the presence or absence of 40µM Pepinh-TRIF (TRIF inhibitor) (n=3 independent experiments). **c**, IL-8 concentration in HepG2 cells treated with OA PA for 24 hours in the presence or absence of 40µM TJ-M2010-5 (MyD88 inhibitor) (n=3 independent experiments). IL-8 concentration in Huh7 cells treated with OA PA for 24 hours in the presence or absence of 5µM (5Z)-7-Oxozeaenol (TAK1 inhibitor) (n=3 independent experiments). **e**, TGFβ1 concentration in LX2 cells following 24 hours treatment with conditioned media from primary hepatocytes treated with OA PA, flagellin in the presence or absence of 1µM TH1020 (TLR5 inhibitor) (n=3 replicates per condition). P values determined by t test or ANOVA with post-test multiple comparisons. ANOVA, analysis of variance. Only statistically significant comparisons (p<0.05) are highlighted.

**Supplementary Table 1.** MASLD participant details for study.

| Age | Gender | BMI | T2DM | Liver category | Fibrosis stage | LS M | Samples used |  |  |  |  |  |
| --- | --- | --- | --- | --- | --- | --- | --- | --- | --- | --- | --- | --- |
|  |  |  |  |  |  |  | Liver qPCR | Stool shotgun metagenomics | LPS ELISA | Flagellin ELISA | HMGB1 ELISA | IL-8 lumine x |
| 39 | F | 24.2 | 0 | Healthy | 0 | 5 |  |  | X |  |  | X |
| 34 | M | 28.7 | 0 | Healthy | 0 |  |  | X | X |  |  | X |
| 39 | M | 44.9 | 0 | Healthy | 0 |  |  |  |  |  |  | X |
| 66 | F | 31.1 | 0 | Healthy | 0 |  |  |  |  |  |  | X |
| 55 | F | 22.9 | 0 | Healthy |  | 3.3 |  |  | X |  |  |  |
| 74 | M | 30.5 | 0 | Healthy |  | 2.8 |  |  | X |  |  |  |
| 27 | M | 26.0 | 0 | Healthy | 0 |  |  |  | X |  |  |  |
| 37 | M | 29.2 | 0 | Healthy | 0 | 5.7 |  | X | X |  |  |  |
| 29 | M | 25.1 | 0 | Healthy | 0 |  |  |  | X |  |  | X |
| 36 | F | 37.4 | 0 | Healthy | 0 |  | X | X | X | X | X |  |
| 56 | F | 44.2 | 0 | Healthy | 0 |  | X | X | X | X | X |  |
| 54 | F | 24.0 | 0 | Healthy | 0 |  |  |  | X |  |  |  |
| 40 | F | 43.5 | 0 | Healthy | 0 |  | X | X | X | X | X |  |
| 64 | M | 22.7 | 0 | Healthy | 0 |  |  |  | X |  |  |  |
| 57 | M | 32.0 | 1 | Healthy | 0 |  | X |  | X | X | X |  |
| 61 | M | 47.6 | 0 | Healthy | 0 |  | X | X | X | X | X |  |
| 35 | M | 21.9 | 0 | Healthy | 0 |  |  |  | X |  |  |  |
| 47 | F | 41.2 | 0 | Healthy | 0 |  |  |  | X |  |  |  |
| 44 | F | 43.1 | 0 | Healthy | 0 |  |  | X | X | X | X |  |
| 37 | F | 31.9 | 0 | Healthy | 0 | 5.8 |  |  |  |  |  |  |
| 46 | F | 43.7 | 0 | Healthy | 0 |  |  | X |  |  | X |  |
| 43 | F | 33.3 | 1 | Healthy | 0 |  | X |  |  | X |  |  |
| 58 | F | 42.0 | 0 | Healthy | 0 |  |  | X | X | X | X |  |
| 59 | F | 29.5 | 0 | Steatosis | 0 |  |  | X | X |  |  | X |
| 66 | F | 31.1 | 0 | Steatosis | 0 |  |  |  | X |  |  |  |
| 44 | F | 38.2 | 0 | Steatosis | 0 |  |  | X | X |  |  | X |
| 55 | M | 42.0 | 1 | Steatosis | 1 |  | X | X | X | X | X |  |
| 62 | F | 35.2 | 1 | Steatosis | 0 |  | X | X | X | X | X |  |
| 43 | F | 45.8 | 1 | Steatosis | 0 |  | X | X | X | X |  |  |
| 36 | F | 44.7 | 0 | Steatosis | 0 |  |  |  | X |  |  |  |
| 52 | F | 43.6 | 0 | Steatosis | 0 |  |  |  | X |  |  |  |
| 38 | F | 45.5 | 0 | Steatosis | 0 |  |  |  | X |  |  |  |
| 34 | F | 47.2 | 1 | Steatosis | 0 |  |  |  | X |  |  |  |
| 60 | F | 30.1 | 1 | Steatosis | 0 |  |  | X |  |  |  |  |
| 52 | F | 38.2 | 0 | Steatosis | 1 |  |  | X |  |  | X |  |
| 51 | F | 35.8 | 1 | Steatosis | 0 |  | X | X |  | X |  |  |
| 28 | M | 27.7 | 0 | Steatosis | 0 | 4 |  |  |  |  |  | X |
| 41 | M | 23.7 | 0 | Steatosis * |  | 7.1 |  |  | X |  |  |  |
| 66 | M | 37.6 | 1 | Steatosis * |  | 5.3 |  | X | X |  |  |  |
| 68 | F | 31.5 | 1 | Steatosis * |  | 5.7 |  |  | X |  |  |  |
| 28 | M | 27.7 | 0 | Steatosis * |  | 4 |  |  | X |  |  |  |
| 52 | M | 29.9 | 0 | Steatosis * |  | 5.4 |  |  | X |  |  |  |
| 64 | M | 28.3 | 0 | Steatosis * |  | 5.7 |  |  | X |  |  |  |
| 59 | M | 32.7 | 1 | Steatosis * |  | 4 |  |  | X |  |  |  |
| 51 | M | 31.7 | 0 | Steatosis * |  | 5.5 |  | X | X |  |  |  |
| 65 | M | 34.9 | 0 | Steatosis * |  | 4.4 |  |  | X |  |  |  |
| 47 | M | 24.7 | 0 | Steatosis * |  | 7.3 |  |  | X |  |  |  |
| 44 | F | 35.5 | 0 | Steatosis * |  | 8.6 |  |  | X |  |  |  |
| 44 | M | 26.9 | 0 | Steatosis * |  | 5 |  | X | X |  |  |  |
| 55 | F | 28.7 | 0 | Steatosis * |  | 5.8 |  | X | X |  |  |  |
| 36 | F | 29.4 | 1 | Steatosis * |  | 5.1 |  | X | X |  |  |  |
| 45 | M | 27.3 | 1 | Steatosis * |  | 6.4 |  |  |  |  |  |  |
| 56 | M | 26.2 | 0 | Steatosis * |  | 9.8 |  | X |  |  |  | X |
| 46 | F | 33.9 | 0 | Steatosis * |  | 4.8 |  |  |  |  |  |  |
| 33 | M | 28.5 | 0 | Steatosis * |  | 4.2 |  |  |  |  |  |  |
| 43 | M | 26.3 | 0 | Steatosis * |  | 6.6 |  | X |  |  |  | X |
| 58 | M | 30.8 | 0 | Steatosis * |  | 7.2 |  | X |  |  |  | X |
| 49 | M | 27.1 | 1 | MASH | 0 |  |  |  | X |  |  |  |
| 50 | M | 36.1 | 1 | MASH | 0 |  |  |  | X |  |  | X |
| 46 | M | 36.6 | 0 | MASH | 1 |  |  |  |  |  |  |  |
| 47 | F | 33.4 | 1 | MASH | 1 |  |  |  |  | X |  | X |
| 35 | F | 35.5 | 1 | MASH | 1 |  |  |  |  |  |  |  |
| 42 | F | 38.8 | 1 | MASH | 1 |  |  |  |  |  |  |  |
| 34 | M | 44.7 | 0 | MASH | 1 |  |  |  |  |  |  |  |
| 50 | F | 39.9 | 1 | MASH | 1 |  |  |  |  |  | X |  |
| 54 | F | 33.1 | 0 | MASH | 1 |  |  |  | X |  |  | X |
| 33 | M | 29.1 | 0 | MASH | 1 |  |  |  | X |  |  |  |
| 52 | M | 24.4 | 0 | MASH | 1 |  |  |  | X |  |  | X |
| 61 | F | 35.4 | 0 | MASH | 1 |  |  | X | X |  |  |  |
| 65 | M | 33.5 | 1 | MASH | 1 |  |  | X | X |  |  | X |
| 29 | F | 30.1 | 1 | MASH | 1 |  |  |  | X |  |  |  |
| 54 | F | 38.3 | 1 | MASH | 1 |  |  | X | X |  |  |  |
| 56 | M | 24.8 | 0 | MASH | 1 |  |  | X | X |  |  |  |
| 49 | F | 24.4 | 0 | MASH | 1 |  |  |  | X |  |  |  |
| 40 | M | 24.9 | 0 | MASH | 1 |  |  | X | X |  |  | X |
| 54 | F | 26.9 | 0 | MASH | 1 |  |  |  | X |  |  | X |
| 36 | F | 43.9 | 0 | MASH | 1 |  | X |  | X | X | X |  |
| 58 | F | 39.8 | 1 | MASH | 1 |  | X | X | X | X |  |  |
| 30 | F | 41.4 | 0 | MASH | 1 |  | X |  | X | X |  |  |
| 40 | F | 53.5 | 0 | MASH | 1 |  |  | X | X | X | X |  |
| 36 | F | 44.7 | 0 | Fibrosis | 1 |  | X |  | X | X |  |  |
| 20 | M | 49.1 | 0 | Fibrosis | 1 |  | X |  | X | X |  |  |
| 47 | M | 27.8 | 0 | MASH | 2 |  |  |  |  |  |  |  |
| 56 | M | 33.8 | 0 | MASH | 2 |  |  |  |  | X |  | X |
| 71 | M | 41.2 | 0 | MASH | 2 |  |  |  | X |  |  | X |
| 66 | F | 22.2 | 0 | MASH | 2 |  |  |  | X |  |  | X |
| 41 | F | 33.0 | 0 | MASH | 2 |  |  |  | X |  |  | X |
| 64 | F | 51.4 | 1 | MASH | 2 |  |  | X | X |  |  | X |
| 55 | M | 32.4 | 0 | MASH | 2 |  |  |  | X |  |  | X |
| 44 | M | 28.4 | 0 | MASH | 2 |  |  |  | X |  |  |  |
| 51 | F | 49.5 | 0 | MASH | 2 |  | X |  | X | X | X |  |
| 53 | M | 48.3 | 1 | MASH | 2 |  | X |  | X | X | X |  |
| 51 | F | 34.8 | 1 | MASH | 2 |  | X | X | X | X | X |  |
| 35 | F | 39.9 | 0 | Fibrosis | 2 |  |  | X |  |  |  |  |
| 62 | F | 24.7 | 0 | MASH | 3 |  |  |  |  | X |  |  |
| 57 | M | 30.7 | 0 | MASH | 3 |  |  |  |  | X |  | X |
| 52 | F | 28.9 | 1 | MASH | 3 |  |  |  |  | X |  |  |
| 51 | M | 32.9 | 1 | MASH | 3 |  |  |  |  |  |  |  |
| 43 | M | 25.6 | 1 | MASH | 3 |  |  |  |  | X |  |  |
| 68 | M | 38.6 | 1 | MASH | 3 |  |  |  |  | X |  |  |
| 66 | F | 23.5 | 1 | MASH | 3 |  |  |  |  | X |  | X |
| 49 | F | 73.6 | 0 | MASH | 3 |  |  |  |  | X |  |  |
| 73 | F | 38.2 | 0 | MASH | 3 |  |  | X |  |  |  |  |
| 68 | M | 37.9 | 1 | MASH | 3 |  |  |  | X |  |  | X |
| 74 | F | 27.7 | 1 | MASH | 3 |  |  | X | X | X |  | X |
| 59 | M | 30.4 | 0 | MASH | 3 |  |  |  | X |  |  |  |
| 36 | M | 24.9 | 1 | MASH | 3 |  |  |  | X |  |  | X |
| 45 | F | 28.1 | 1 | MASH | 3 |  |  | X | X | X |  | X |
| 35 | M | 31.8 | 1 | MASH | 3 |  |  | X | X | X |  |  |
| 60 | M | 44.2 | 0 | MASH | 3 |  |  | X | X |  |  |  |
| 51 | F | 30.9 | 1 | MASH | 3 |  |  | X | X | X |  | X |
| 63 | M | 30.0 | 1 | MASH | 3 |  |  | X | X | X |  | X |
| 49 | F | 27.8 | 1 | MASH | 3 |  |  | X | X |  |  |  |
| 58 | F | 50.1 | 1 | MASH | 3 |  | X | X | X | X | X |  |
| 35 | F | 51.5 | 0 | MASH | 3 |  | X | X | X | X | X |  |
| 63 | F | 57.3 | 1 | MASH | 3 |  | X |  | X | X |  |  |
| 54 | F | 47.4 | 0 | MASH | 3 |  |  | X | X | X | X |  |
| 38 | F | 58.5 | 1 | MASH | 3 |  |  |  | X |  |  |  |
| 51 | M | 26.1 | 1 | MASH | 3 |  |  |  |  |  |  | X |
| 37 | M | 28.9 | 0 | MASH | 3 |  |  |  |  |  |  |  |
| 74 | M | 41.3 | 1 | MASH | 3 |  |  |  |  |  |  | X |
| 66 | F | 43.6 | 0 | Fibrosis | 3 |  |  | X |  |  |  |  |
| 56 | M | 34.2 | 1 | Cirrhosis | 4 |  |  |  | X | X |  |  |
| 66 | M | 30.1 | 1 | Cirrhosis | 4 |  |  | X | X |  |  | X |
| 54 | M | 27.9 | 1 | Cirrhosis | 4 |  |  |  |  | X |  |  |
| 55 | M | 25.9 | 1 | Cirrhosis | 4 |  |  |  |  | X |  |  |
| 51 | M | 38.1 | 1 | Cirrhosis | 4 |  |  |  | X | X |  |  |
| 59 | F | 33.9 | 1 | Cirrhosis * |  | 21.3 |  |  | X | X |  |  |
| 65 | M | 31.1 | 0 | Cirrhosis * |  | 31.5 |  | X | X |  |  |  |
| 73 | M | 28.8 | 1 | Cirrhosis * |  | 14.5 |  | X | X | X |  |  |
| 67 | M | 39.1 | 1 | Cirrhosis * |  | 10.2 |  |  |  | X |  |  |
| 31 | M | 58.1 | 0 | Cirrhosis * |  | 60 |  |  | X |  |  |  |
| 76 | F | 34.1 | 1 | Cirrhosis * |  | 20.7 |  | X | X | X |  |  |
| 67 | M | 29.7 | 1 | Cirrhosis * |  | 11.8 |  |  |  | X |  |  |
| 49 | M | 33.4 | 0 | Cirrhosis * |  | 10.3 |  |  | X |  |  |  |
| 73 | F | 34.3 | 0 | Cirrhosis * |  | 21.3 |  | X | X |  |  |  |
| 74 | F | 37.1 | 1 | Cirrhosis * |  | 38.7 |  | X | X |  |  | X |
| 58 | M | 45.9 | 1 | Cirrhosis * |  | 36.2 |  | X | X |  |  |  |
| 71 | M | 27.8 | 1 | Cirrhosis * |  | 45.4 |  | X | X |  |  |  |
| 74 | M | 35.9 | 1 | Cirrhosis * |  | 8.9 |  |  |  | X |  |  |
| 60 | M | 32.1 | 0 | Cirrhosis * |  | 10.1 |  |  |  | X |  |  |
| 63 | M | 39.1 | 1 | Cirrhosis * |  | 32.8 |  |  |  | X |  | X |

Gender: M – male, F Female, BMI – body mass index, T2DM – Type 2 diabetes mellitus, 0 – no type 2 diabetes mellitus, 1 – known diagnosis of type 2 diabetes mellitus, LSM – Liver stiffness measurement in kPa, \* - liver disease diagnosed through radiology, not by histology. X indicates sample used

**Supplementary Table 2.** Hepatic TLR2, TLR4, TLR5 expression in independent MASLD datasets Combined UCAM and VCU cohort.

| **Gene** | **L2FC_UC**<br>**AM/VCU:**<br>**Mild vs**<br>**Control** | **P-**<br>**VALUE_U**<br>**CAM/VCU:**<br>**Mild vs**<br>**Control** | **P-**<br>**ADJUSTED_**<br>**UCAM/VCU:**<br>**Mild vs**<br>**Control** | **L2FC_UC**<br>**AM/VCU:**<br>**Moderate**<br>**vs Mild** | **P-**<br>**VALUE_U**<br>**CAM/VCU:**<br>**Moderate**<br>**vs Mild** | **P-**<br>**ADJUSTED_**<br>**UCAM/VCU:**<br>**Moderate vs**<br>**Mild** | **L2FC_UC**<br>**AM/VCU:**<br>**Severe vs**<br>**Mild** | **P-**<br>**VALUE_U**<br>**CAM/VCU:**<br>**Severe vs**<br>**Mild** | **P-**<br>**ADJUSTED_**<br>**UCAM/VCU:**<br>**Severe vs**<br>**Mild** |
| --- | --- | --- | --- | --- | --- | --- | --- | --- | --- |
| **TLR5** | -0.191 | 0.321 | 0.612 | 0.086 | 0.320 | 0.648 | 0.365 | 0.009 | **0.048** |
| **TLR4** | 0.045 | 0.776 | 0.907 | 0.049 | 0.461 | 0.754 | 0.040 | 0.642 | 0.783 |
| **TLR2** | 0.366 | 0.032 | 0.184 | 0.063 | 0.396 | 0.707 | 0.128 | 0.311 | 0.502 |

**EPoS cohort**
| **Gene** | **L2FC_EPoS:**<br>**Moderate vs**<br>**Mild** | **P-VALUE_EPoS:**<br>**Moderate vs Mild** | **P-**<br>**ADJUSTED_EPoS:**<br>**Moderate vs Mild** | **L2FC_EPoS:**<br>**Severe vs**<br>**Mild** | **P-**<br>**VALUE_EPoS:**<br>**Severe vs Mild** | **P-**<br>**ADJUSTED_EPoS:**<br>**Severe vs Mild** |
| --- | --- | --- | --- | --- | --- | --- |
| **TLR5** | 0.246 | 0.030 | 0.511 | 0.677 | 0.000 | **0.000** |
| **TLR4** | 0.081 | 0.496 | 0.916 | -0.024 | 0.882 | 0.956 |
| **TLR2** | 0.094 | 0.351 | 0.868 | 0.210 | 0.115 | 0.336 |

Mild = MASLD with F0; Moderate = MASLD with F1-2; Severe = MASLD with F3-4

**Supplementary Table 3.** Chronic hepatitis B participant details for study.

| **Age** | **Gender** | **BMI** | **LSM** |
| --- | --- | --- | --- |
| 31 | F | 24.0 | 3.7 |
| 29 | F | 18.7 | 6.3 |
| 35 | M | 23.6 | 8.2 |
| 33 | M | 25.9 | 7.2 |
| 63 | M | 26.7 | 7.8 |
| 47 | M | 25.2 | 2.5 |
| 25 | M | 24.0 | 3.9 |
| 24 | M | 24.1 | 8.2 |
| 60 | M | 28.5 | 3.7 |
| 42 | M | 23.1 | 12.7 |
| 26 | M | 22.7 | 7.8 |
| 21 | M | 26.8 | 4.4 |
| 23 | F | 23.5 | 4.0 |
| 66 | M | 24.5 | 7.4 |
| 18 | M | 18.7 | 13.9 |
| 23 | M | 21.4 | 5.0 |
| 28 | M | 23.0 | 35.1 |
| 47 | M | 26.1 | 20.2 |

Gender: M – male, F Female, BMI – body mass index, LSM – Liver stiffness measurement in kPa

**Supplementary Table 4.** Inhibitors, concentrations and treatment protocol for in vitro experiments.

| Inhibitor | Concentration | Treatment protocol |
| --- | --- | --- |
| TH1020 | 1µM | added 18 hours prior to treatment of cells |
| PAb hTLR4 | 5µg/mL | added 15 minutes prior to treatment of cells |
| PAb hTLR2 | 5µg/mL | added 15 minutes prior to treatment of cells |
| Obeticholic acid | 10µM (primary hepatocytes),<br>2µM (HepG2, Huh7) | added 18 hours prior to treatment of cells |
| SML2148 | 80µM primary hepatocytes),<br>16µM (HepG2, Huh7) | added 18 hours prior to treatment of cells |
| Lipofermata | 50µM (primary hepatocytes),<br>10µM (HepG2, Huh7) | added 18 hours prior to treatment of cells |
| Pepinh-TRIF | 40µM | added 18 hours prior to treatment of cells |
| TJ-M2010-5 | 40µM | added 18 hours prior to treatment of cells |
| CU-CPT4a | 27µM | added 18 hours prior to treatment of cells |
| (5Z)-7-Oxozeaenol | 5µM | added 18 hours prior to treatment of cells |

TH1020 – TLR5 inhibitor, PAb hTLR4 – TLR4 inhibitor, PAb hTLR2 – TLR2 inhibitor, Obeticholic acid – FATP5 inhibitor, SML2148 – CD36 inhibitor, Lipofermata – FATP2 inhibitor, Pepinh-TRIF – TRIF inhibitor, TJ-M2010-5 – MyD88 inhibitor, CU-CPT4a – TLR3 inhibitor, (5Z)-7-Oxozeaenol – TAK1 inhibitor

**Supplementary Table 5.** List of ELISA kits used in this study.

| Item | Supplier | Product # |
| --- | --- | --- |
| Human Flagellin ELISA | MyBioSource | MBS268117 |
| Human Lipopolysaccharide ELISA | MyBioSource | MBS266722 |
| Human IL-8 ELISA | R&D systems | DY208 |
| Human Pro-collagen 1α1 ELISA | R&D systems | DY6220 |
| Human TGFβ1 ELISA | R&D systems | DY240 |
| Human HMGB1 ELISA | Novus Biologicals | NBP2-62766 |
| Human DDIT3 ELISA | Abcam | ab315074 |
| Human Fatty Acid Oxidation In-Cell ELISA | Abcam | ab118182 |

**Supplementary Table 6.**
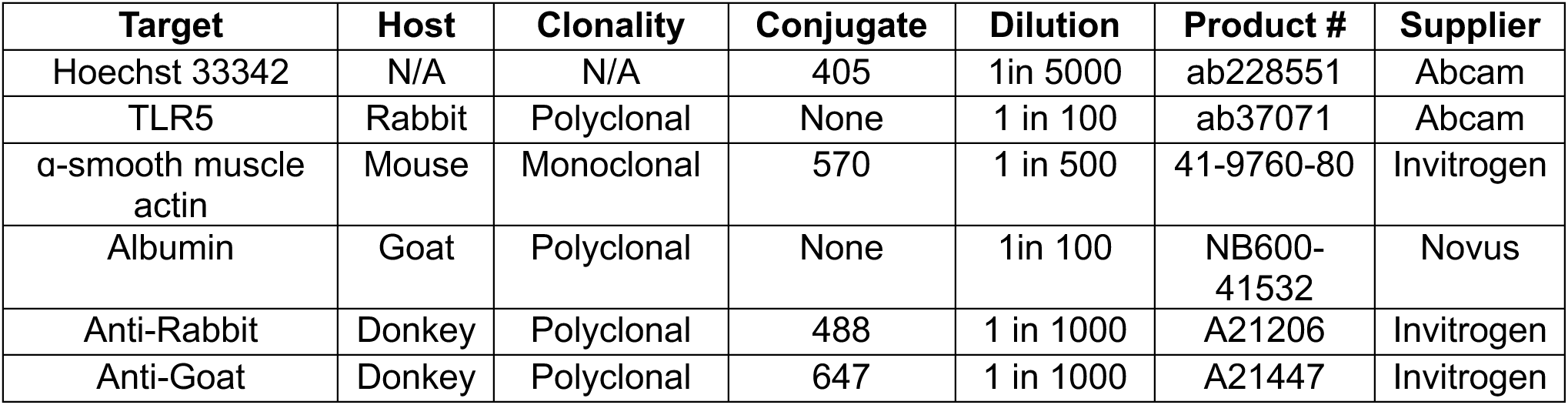
List of antibodies used for immunofluorescence.

## Acknowledgements

We are grateful to Salma Samsuddin, Victoria Berryman, Johanna Preston, Melanie Pattrick, Caroline Sutcliffe and Noorshad Joti for facilitating patient recruitment for this study.

## Author contributions

Conceptualisation: WL, WA

Methodology: WL, NNW, IGM, CM, JB, RW, ND, RG, JL, KD, HM, AG, TJK, JF, MICM, KR, MV, DEA, SA, OG, WA

Investigation: WL, NNW, RK, IGM, UG, JB, CM, JB, RW, ND, RG, MJR, TJK, MICM, KR, MV, DEA, SA, OG

Visualisation: WL, JB, RW, MJR, TJK, MICM, DEA, SA, OG

Funding acquisition: WL, WA Project administration: WL, GH, WA Supervision: WA

Writing (original draft): WL

Writing (review and editing): WL, UG, ND, TJK, JF, KR, MV, DEA, SA, OG, WA

## Data transparency

Data, analytic methods can be requested via direct contact with corresponding author

